# Evidence for adaptive morphological plasticity in the Caribbean coral, *Acropora cervicornis*

**DOI:** 10.1101/2022.03.04.483038

**Authors:** Wyatt C. Million, Maria Ruggeri, Sibelle O’Donnell, Erich Bartels, Cory J. Krediet, Carly D. Kenkel

## Abstract

Genotype-by-environment interactions (GxE) indicate that variation in organismal traits cannot be explained by fixed effects of genetics or site-specific plastic responses alone. For tropical coral reefs experiencing dramatic environmental change, identifying the contributions of genotype, environment, and GxE on coral performance will be vital for both predicting persistence and developing restoration strategies. We quantified the impacts of G, E, and GxE on the morphology and survival of the endangered coral, *A. cervicornis*, through an in situ transplant experiment exposing common garden (nursery) raised clones of ten genotypes to nine reef sites in the Florida Keys. By fate-tracking outplants over one year with colony-level 3D photogrammetry, we uncovered significant GxE on coral size and survivorship indicating that no universal winner exists in terms of colony performance. Moreover, the presence of GxE also implies the existence of intraspecific variation in phenotypic plasticity. Rather than differences in mean trait values, we find that individual-level morphological plasticity is adaptive in that the most plastic individuals also exhibited the fastest growth and highest survival. This indicates that adaptive morphological plasticity may continue to evolve, influencing the success of *A. cervicornis* and resulting reef communities in a changing climate. As focal reefs are active restoration sites, the knowledge that variation in phenotype is an important predictor of performance can be directly applied to restoration planning. Taken together, these results establish *A. cervicornis* as a system for studying the eco-evolutionary dynamics of phenotypic plasticity that also can inform genetic- and environment-based strategies for coral restoration.

## Introduction

Intraspecific variation in phenotype provides raw material for selection to act on resulting in the evolution of trait means (1). However, trait values may also change as individuals are exposed to different environments via phenotypic plasticity (2). While plastic trait changes typically occur within a generation, they have the ability to alter fitness-related traits and promote acclimation, and are therefore relevant for populations experiencing new or stressful environmental conditions (3–6). Moreover, variation in the degree of plasticity can magnify differences among individuals. In the light of intraspecific variation in plasticity, the evolution of trait means becomes dependent not only on individual trait values but also the environments those individuals face. Long-standing theory supports a role for plasticity in trait evolution (7–9) and the presence of significant intraspecific variation in plasticity, i.e. genotype-by-environment interactions (GxE), suggest that plasticity itself can also evolve (10–12).

The evolution of phenotypic plasticity, and consequently its ecological impacts, can occur if variation in plasticity among individuals results in variation in fitness. Selection is expected to increase plasticity when the benefits of producing an environment-specific phenotype outweigh the fitness consequences arising from either the cost of altering that phenotype, or from the production of an underdeveloped phenotype relative to locally adapted individuals (13, 14). Evolutionary models suggest plasticity will be favored in species with high dispersal that will experience predictably high spatial or temporal environmental variation and when the costs of plasticity are low (14). However, limited empirical tests for an adaptive role of plasticity (12, 14) provide inconsistent support for model predictions with variation evident among traits, species, and environments (14–16). More experiments that quantify the fitness costs or benefits of plasticity, especially in nonmodel systems, will improve our broad understanding of it’s ecological and evolutionary role (2, 10) while also uncovering system-specific functions contributing to acclimation to environmental change. This will be particularly important for species of conservation concern, where persistence may be reliant on both adaptive plasticity and the ability of human interventions to leverage it.

Reef-building corals form the base of the most biodiverse marine ecosystems, tropical coral reefs (17, 18). The ecological and economic services they provide are determined in part by the complex three-dimensional structures created by the corals themselves (19–21). This structure provides habitat space (22), reduces wave energy (21), and sustains biological diversity and productivity (19, 23) which support a multi-billion-dollar tourism industry (24). However, these ecosystem services are being lost as wild populations decline due to natural and anthropogenic factors (25, 26). For example, populations of *Acropora cervicornis*, one of two branching coral species once dominating Caribbean reefs, have declined precipitously since the 1970s (27) contributing to a loss of structural complexity (28, 29). This decline has prompted a global effort to understand factors that promote coral survival and persistence under changing ocean conditions. Both differences in fitness-related traits among common-gardened genotypes and of clones under different conditions (30–33) suggests that some individuals or environments could be used to reestablish the structure and function of reefs (34–36). However, as corals experience new conditions, via translocation during reef restoration (37, 38) or through climate change (39, 40), it is unclear whether top performing genotypes will maintain their status (41, 42) or if variation in plasticity will result in new ‘winners’ and ‘losers’ (43). Therefore, clarifying the role of both the environment and genetic background on coral performance as well as GxE will be critical for leveraging naturally occurring biological variation for the restoration of degraded reefs.

Phenotypic plasticity has been commonly documented in coral morphology (44, 45), physiology (46, 47), and gene expression (48, 49) in response to a variety of abiotic factors, indicating the environmental responsiveness of phenotypes, some of which is correlated with fitness-related traits at the population level (50–52). Within-populations, genotype-by-environment interactions have been reported less frequently (41, 43, 53) but the relationship between phenotypic plasticity and overall fitness remains unresolved. This gap in knowledge limits both our understanding of the evolutionary potential of plasticity and its role in the natural and human-assisted recovery of coral reefs in a changing climate.

While variation in plasticity exists in wild coral populations (41, 53, 54), it is unclear if such variation is maintained in restoration corals, like *A. cervicornis*, that have been propagated in common-garden nurseries for decades. Given that restoration involves outplanting clonal replicates (ramets) across environmentally-diverse, natural reefs (37, 55), understanding how individual phenotypes vary can help direct restoration efforts. Previous efforts found no evidence of GxE in *A. cervicornis* morphology despite strong site and genotype effects on linear growth and survival (33, 37, 55) but significant variation in bleaching responses among genotypes was observed across natural reefs (43). GxE in fitness-related traits, like bleaching tolerance, suggest that a multisite phenotype may more accurately describe *A. cervicornis* performance in nature.

We rigorously fate-tracked 270 restored coral colonies on natural reefs in a multi-site transplant experiment to test the effect of genotype, environment and their interaction on growth rate, size, risk of fragmentation, and survival (Fig. 1A-C). We uncovered significant GxE in survival and absolute size, measured with 3D photogrammetry, and identified relationships between morphological plasticity, growth, and survival that support the presence of adaptive plasticity in *A. cervicornis*. Taken together, our results establish a new system in which to investigate the ecological and evolutionary impacts of adaptive plasticity while also informing reef restoration.

**Figure 1:**
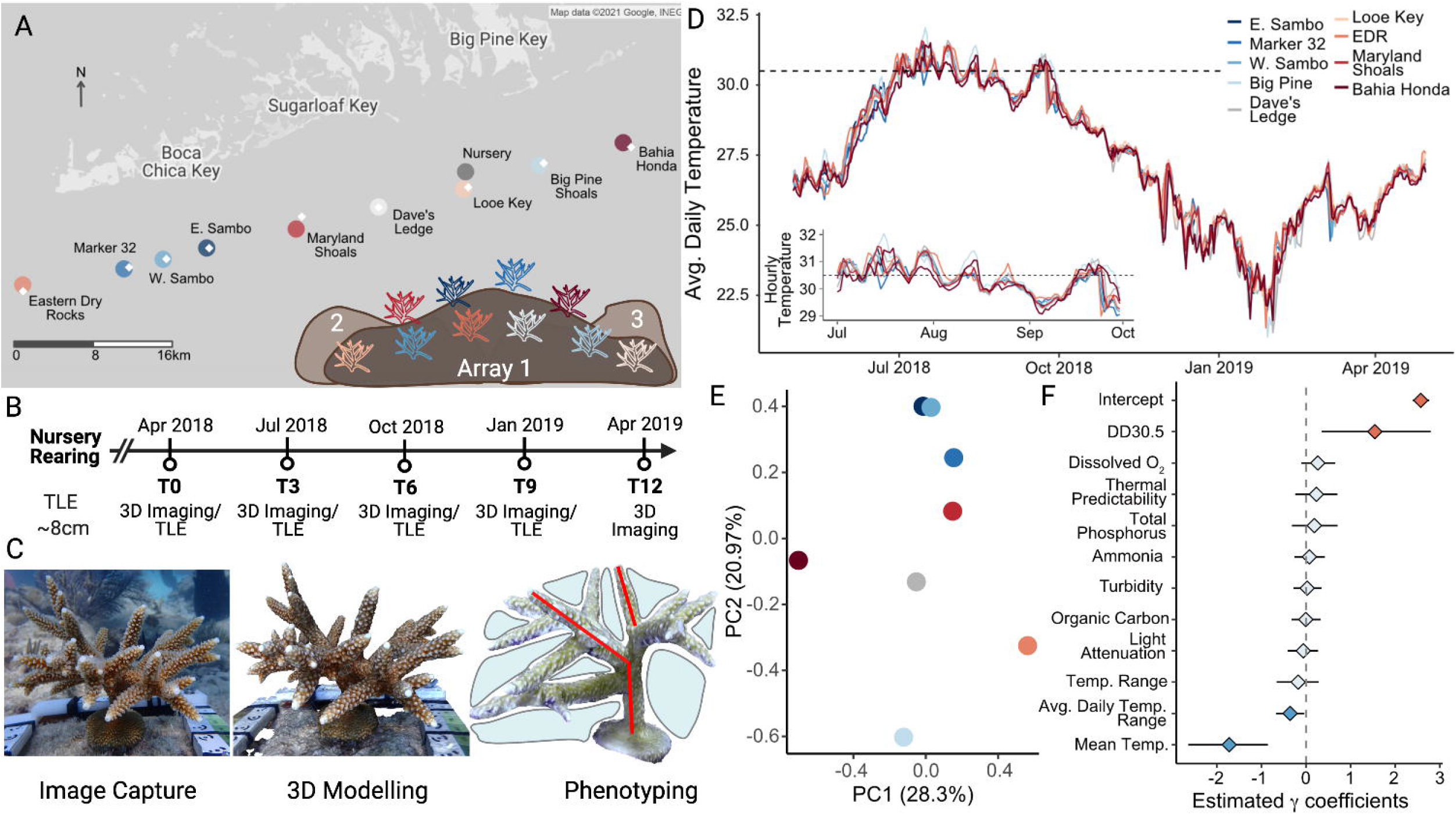
Experimental design and environmental conditions **A)** Location of outplant sites (Table S1) and the source restoration nursery in the lower Florida Keys, USA. Ten genets (genotypes) of *A. cervicornis* were outplanted in triplicate to nine reef sites in three sub-arrays. Sites are colored by average survival from dark blue (highest) to dark red (lowest) and the location of site-specific SERC water quality monitoring stations (56) are indicated by the white diamonds. **B)** Ramets were reared in the nursery to a mean size of 8 cm and were measured just prior to outplanting in April 2018 **C)** To obtain growth and morphology data, still images were captured using underwater photography, which were used to build 3D models in Agisoft Metashape, that were subsequently measured for total linear extension (TLE; example red lines), surface area (SA), volume (V), and volume of the interstitial space (V_inter_; example shaded blue area) **D)** Average daily temperature of each site (colored by survival rank) for the one-year experimental period. Inset shows hourly temperatures from July through September 2018. The dashed line indicates the local bleaching threshold of 30.5°C. **E)** Principal components analysis of historical SERC environmental metrics characterizing outplant reef sites (Looe Key is excluded due to missing data) **F)** Results of a Bayesian negative binomial generalized linear mixed effects model testing the association of eleven uncorrelated environmental parameters on the change in V_inter_. Horizontal black lines indicate 95% credible intervals of the posterior distributions. Values above (red) or below (blue) indicate significant association between the variable and the change in V_inter_ across sites.

## Results

### Ramet survival is a function of genotype, outplant site, and the interaction

Of the 270 outplanted ramets, 48 died and 24 were declared missing leaving 198 surviving ramets at the end of one year. Cox proportional hazard models showed significant effects of genotype (p=0.002) and site (p=0.005) on survival. The model including the interaction term, while improving fit, did not converge, so the additive model was used to obtain risk scores for each genotype and site (Table S2-3). On average, Genotype (G) 36 had the highest ramet survival (96%) while G41 had the lowest (61.5%) across sites, which incurred a mortality risk 2.6 times that of G36 on average (Fig. 2A, Table S2). Three genotypes (G41, G62, G13) form a group of high risk genets (>2.3x higher mortality risk in comparison to G36) while remaining genets display intermediate risk ranging from nearly equivalent to 1.7 times that of G36 (Fig. 2A, Table S2). Ramets outplanted to Bahia Honda (60% survival) had a 2.5 times greater risk of mortality than those outplanted to E. Sambo, the site with the highest survival on average (96.4%, Fig. 2B). Similar to average genotype scores, sites exhibited a continuum of increasing risk with mortality ranging from 1.1 to 2.5 times higher than the reference, E. Sambo, with the highest mortality risk occurring at Bahia Honda (Table S3).

**Figure 2:**
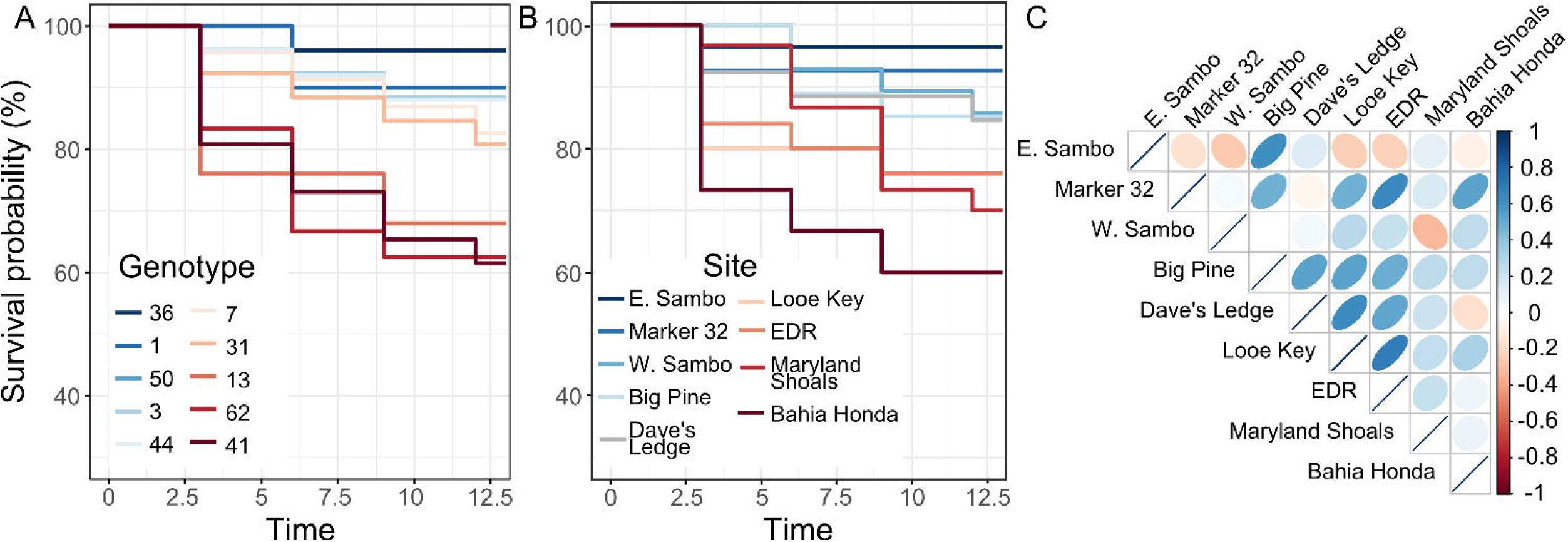
Genotype, environment, and GxE patterns of survival. **A)** Survival probability for genets and **(B)** sites over the one-year experimental period. Genets and sites are colored by decreasing overall survival from blue to red. **C)** Pairwise correlations of genet survival rank across outplant sites. Ellipse shape and color is proportional to the strength and direction of the correlation between two sites. Sites are ordered according to survival as in (B).

Pairwise correlations of genet survival ranks across sites show the identity of the best surviving genet is not maintained across sites, with most correlations being close to 0 (Fig. 3C). The highest positive correlations (Pearson’s correlation = 0.54 to 0.70) were observed among sites with intermediate survival on average (Big Pine, Dave’s Ledge, Looe Key, and EDR, Fig. 2B) and not necessarily among geographically neighboring sites (Fig. 1A, S1).

**Figure 3:**
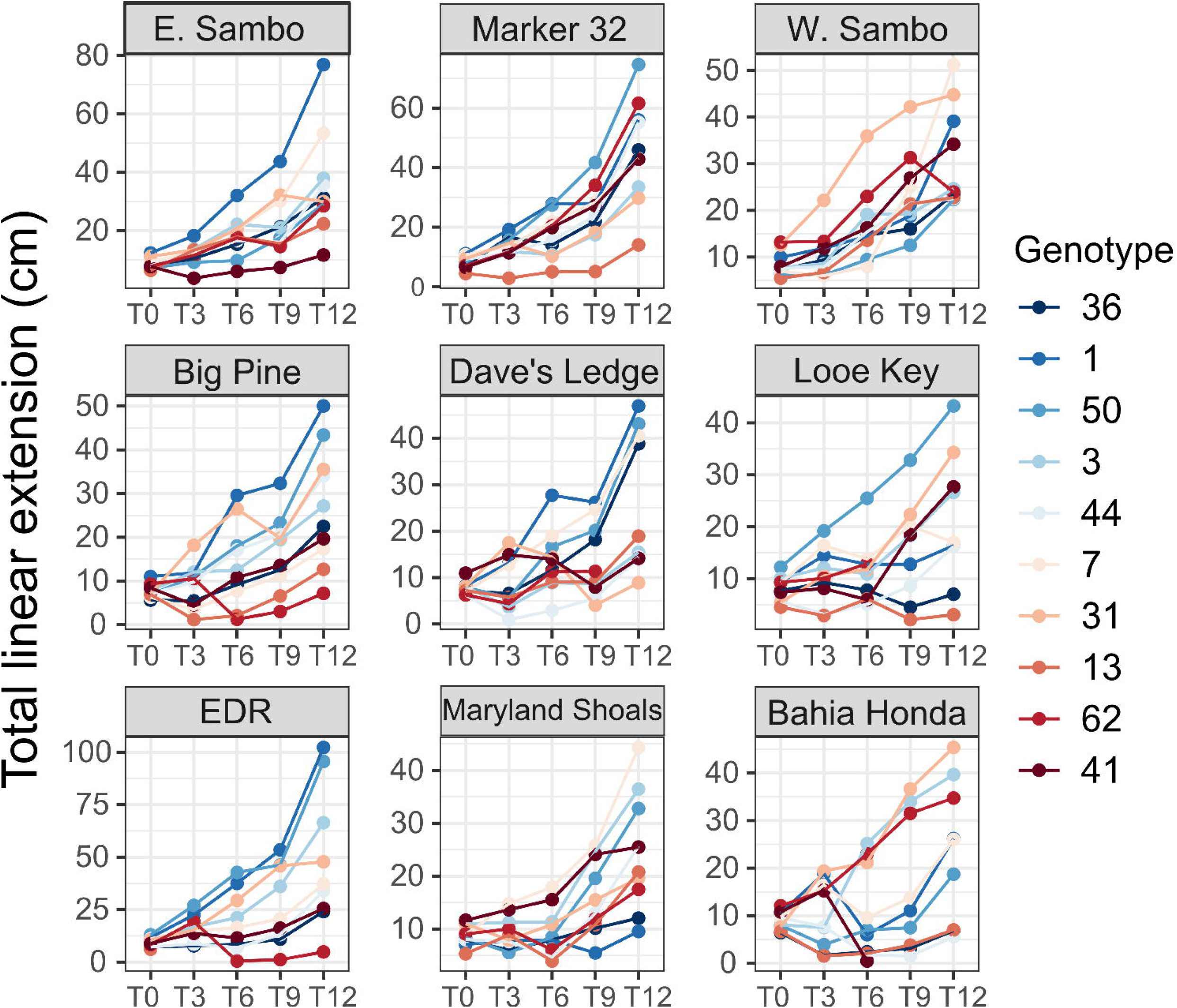
Average size in total linear extension (TLE) for each genotype (colored by survival probability as in Fig. 2A) over time. Reef sites are also ordered by survival probability (left to right).

### Non-random fragmentation in *A. cervicornis*

We documented 177 instances of fragmentation throughout this experiment. Most events occurred within the first three months post-outplant (84) followed by continually decreasing occurrences in subsequent time periods. Cumulative linked models show a significant effect of genotype and site on the likelihood of fragmentation (Table S4) with G1and Marker 32 experiencing the least amount of breakage among genotypes and sites, respectively (Fig. S2). G44 and Bahia Honda experienced the most breakage among genotypes and sites, respectively, during the outplant period. Of the 177 fragmentation events, only 15.8% occurred in the same time period that the ramet died, indicating that breakage did not result in immediate mortality. Fragmentation was not overly prevalent in larger size class colonies, and instead was more common in colonies less than 5 cm and between 5-10 cm in length according to Fisher Exact tests (Table S5).

### Morphology exhibits plasticity that varies by genotype (GxE)

We assessed the fixed effects of genotype, outplant site, time, their interactions, as well as initial size on the absolute size in four traits: total linear extension (TLE), surface area (SA), volume (V), and volume of interstitial space (V_inter_) (Table S6). Ramets experiencing fragmentation were retained, and the site-specific array to which the ramet was outplanted and the number of fragmentation events it experienced were included as random effects (Table S7). Significant genotype-by-site interactions were detected in all four traits (p<0.001), whereas no effects were detected for the genotype-by-time and genotype-by-site-by-time interactions in any trait (p>0.05). Absolute size trajectories varied among genets and inconsistent genet rank order was evident across sites, confirming the existence of GxE (Fig. 3, Fig. S3-5). Significant effects of genotype, time, and initial ramet size were also evident for all traits (p<0.0001). Absolute trait size increased over time and with the initial size of a ramet (Fig. 3, S3-5). Ramets of G50 were the largest on average, while G13’s ramets were the smallest. The largest ramet sizes were reached at EDR, a site with the third worst survival on average (Fig. 2B). However, significant fixed effects of outplant site were not evident, while significant site-by-time interactions were found in SA and V (p<0.01). Random effects of fragmentation (p<0.0001) and array within site (p<0.0001) were also evident in all trait models (Table S7).

### Growth rate is dependent on genotypic and environmental characteristics

Growth rate (unit/month) in each trait (TLE, SA, V, V_inter_), was modeled as a fixed effect of genotype, outplant site, time, and their interactions. Size at the beginning of each time interval was included as a fixed effect in linear mixed models to account for potential size-specific growth rates (Table S8). To accurately quantify growth rate, time periods where a ramet experienced negative growth due to fragmentation were excluded from the mixed models, following (33). Fragmentation and the site-specific array to which the ramet was outplanted were included as random effects (Table S9) and despite being removed from the analysis, a moderate effect of breakage persisted for V (p=0.11). Genotype-by-site and the 3-way interaction between genotype, site, and time were not significant for the growth rate of any traits despite the significant GxE in absolute size. Significant genotype-by-time and site-by-time interactions were detected in TLE, SA, V_inter_ (p<0.01) and TLE, V, and V_inter_ (p<0.01), respectively. A significant fixed effect of genotype was present in TLE, V, and V_inter_ (p<0.05). Growth rate in all traits exhibited a significant fixed effect of time (p<0.0001). Growth rate increased with increasing initial size and time in all traits (p<0.0001, Fig. S6) but standardized growth, calculated as growth rate per unit of existing tissue following (57), decreased with size (Fig. S7). Fixed effects of outplant site were only evident in V (p=0.03) while variation among arrays within sites was nonsignificant in V and SA despite significant array effects on TLE and V_inter_ (p<0.05).

### The capacity for morphological plasticity is correlated with improved survival and growth

Genotype-by-site interactions in the absolute size of all traits indicates significant variation in the capacity for morphological plasticity among genets. We quantified plasticity using a joint regression analysis (58) which integrates trait data across multiple sites to provide a genotype-specific value of plasticity relative to the population. We found consistent correlations between the degree of trait plasticity, overall mortality risk, and mean growth rate that support an adaptive role (Fig. 4A). Plasticity in absolute size in TLE, SA, V, and V_inter_ was only weakly associated with mortality risk at T3 (R=-0.33 to 0.14, p=0.14 to 0.87, Fig. S8). However, negative relationships, indicating reduced mortality risk in more plastic genotypes, strengthened over time with significant relationships in SA and V evident after 9 months (R = −0.65, p = 0.042 and R = −0.82, p = 0.004, respectively), and in TLE, SA, and V after 12 months (R = −0.69, p = 0.029; R = −0.71, p = 0.02; and R = −0.67, p = 0.033; Fig. S8). Similarly, relationships between average growth rate and trait plasticity began as neutral/weakly positive and progressed to strong positive correlations over time, with the strongest correlations found after 12 months (R= 0.81 to 0.85, p= 0.0019 to 0.0043, Fig. S9). Growth rate tended to be negatively correlated with mortality risk, indicating that higher growth rates were associated with decreased mortality risk, although significant relationships were only observed at later time points (V, 9 months, R= −0.7, p=0.023; TLE, 12 mos, R=-0.66, p=0.037; SA, 12 mos, R=-0.66, p=0.039, Fig. S10).

**Figure 4.**
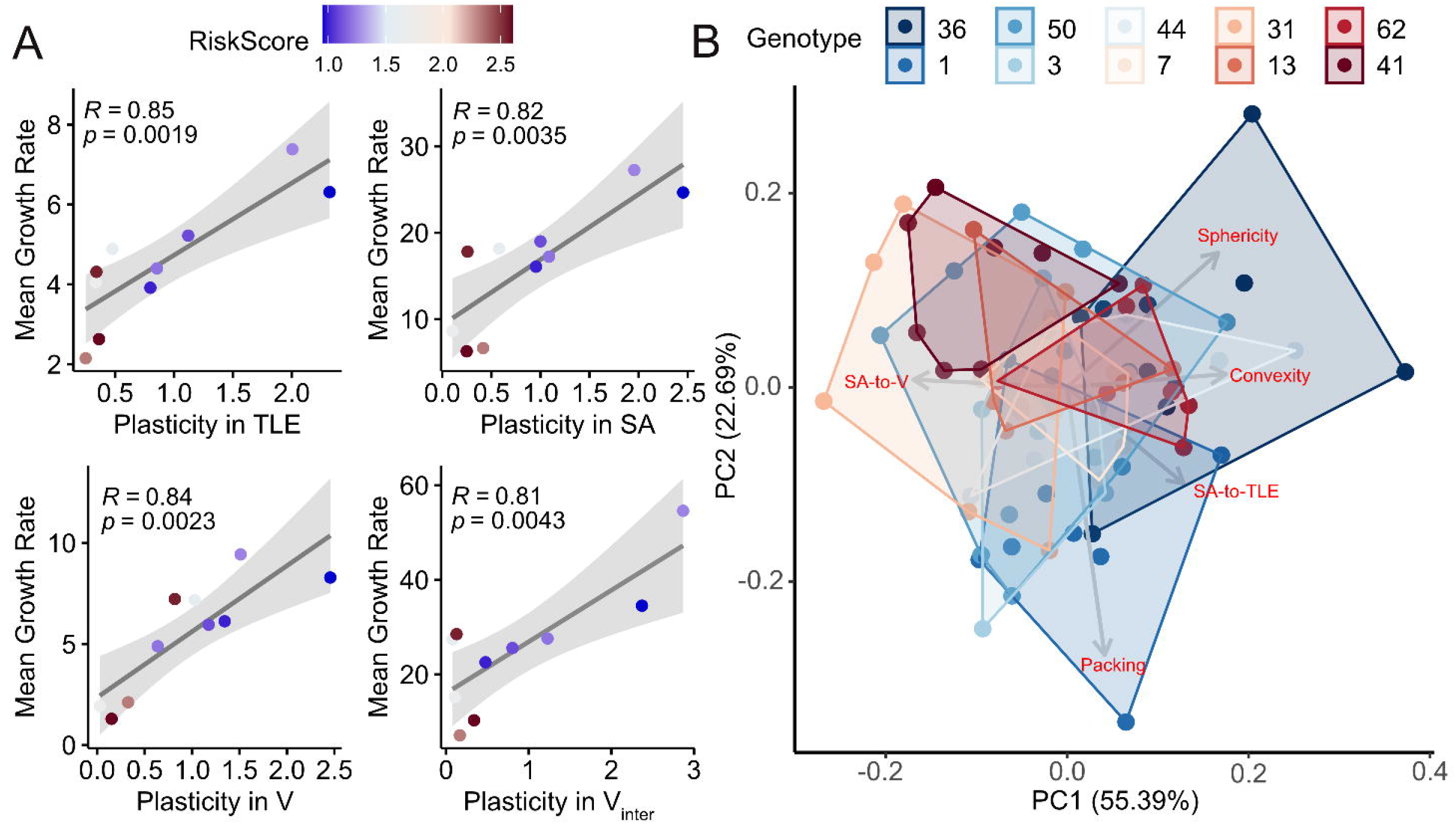
Morphological plasticity and its relationship to growth and survival. **A)** Relationship between genet plasticity in absolute size and average genet growth rate over the final 3 months of the outplant period. Points are colored by genet mortality risk score. Line and shaded region show line of best fit and 95% confidence interval for each relationship. **B)** Principal components analysis of size-independent morphological traits: sphericity, convexity, packing (59), and SA-to-V ratio and SA-to-TLE ratio (gray vectors labeled in red). Points represent individual ramets colored by genet identity (n=5-12 ramets per genet) by decreasing survival from blue to red. Shaded regions (colored by survival rank) frame the most extraneous ramets for each genet and outline the morphospace occupied by a genet.

As a way to assess changes in colony shape regardless of size, we calculated SA-to-V ratio, TLE-to-V ratio, packing, convexity, and sphericity (59) for ramets surviving to T12 with no fragmentation. This subset represents ramets occupying a morpho-space enabling survival and growth uninterrupted by breakage. When plotted in multivariate trait space, ramets did not cluster by genet or site (Fig. 4B, S11). However, ramets of genets showing higher average survival occupy a broader area, or trait space, compared to genets with lower survival (Fig. 4B).

### Variation among offshore reefs may contribute to site-specific growth and survival

Benthic temperatures were recorded hourly at all sites for the one-year experimental period except for Looe Key where data from April 2018 to October 2018 are missing due to a flooded logger (Fig. S12). Therefore, the temperature profiles of 8 sites were used to analyze thermal differences among reefs. Water temperatures over the experimental period were similar between sites (Fig. 1D, Table S10). Annual mean temperatures varied by 0.213°C between the warmest (EDR) and coolest (Bahia Honda) sites. Annual temperature ranges varied from 10.12°C (E. Sambo) to 12.26°C (Bahia Honda) while average daily temperature ranges varied from 0.59°C (Marker 32) to 0.71°C (Bahia Honda). Big Pine experienced the most days where the temperature was at or above 30.5°C (69) while Bahia Honda experienced the fewest (54). Interestingly, Big Pine and Bahia Honda both had the most days above 32°C (3) while the majority of sites never reached this temperature throughout the entire experimental period. Summer thermal predictability, calculated as the sum of positive temporal autocorrelation from July through September, was highest at W. Sambo and lowest at Bahia Honda. The three sites with the highest survival probability had the three highest thermal predictability values.

Site-specific biogeochemical parameters, obtained from the long-term SERC water quality monitoring program (56), did not differ when restricting the dataset to the experimental period. However, analysis of the full historical dataset (1995-2019) revealed significant differences in nitrate and silica dioxide concentrations among the 9 reef sites (p<0.05, Fig S11). A principal components analysis of all thermal and water quality parameters showed large aggregate differences between sites despite this limited variation in individual parameters (Fig. 1E, S13, S14). Sites with the highest survival (E. Sambo, Marker 32, and W. Sambo) clustered together while the remaining sites were broadly distributed. These high survival sites were also associated with high thermal predictability and historical high average light attenuation (Fig. S14). Bahia Honda, the site with the lowest survival and growth, consistently stood out as the most extreme point (Fig. 1E, S13, S14).

After removing highly correlated water quality metrics, eleven parameters were used as candidate variables in a Bayesian negative binomial generalized linear model to assess their power to predict changes in coral morphology and mortality risk. Average temperature was significantly negatively associated with the change in TLE, SA, V, and V_inter_ (Fig. 1F, Fig. S15-18) while average daily temperature range was negatively associated with only V_inter_. Days above 30°C was significantly positively associated with change in V_inter_ but no patterns were evident for other traits (Fig. S15-18). Risk score was not significantly associated with any of the environmental parameters (Fig. S19).

## Discussion

The presence of a significant GxE interaction indicates that individuals differ in their sensitivity to environmental variation. This variation in reaction norm slope among individuals can reduce prediction accuracy and confound selective breeding programs (60). Here, we identify significant GxE in both the size and ultimate survival of restoration lines of *A. cervicornis* indicating that no single genet ‘wins’ in all contexts when considering the change in mean trait values across environments. However, the existence of GxE also means that there is genetic variation in the capacity for plasticity, or the degree of environmental responsiveness of individual genotypes. Rather than mean size, we find that this plasticity, or the degree to which a genet is able to change its size relative to the population mean across sites, is positively correlated with mean growth rate and survival. This suggests that plasticity may continue to evolve, although context-dependent trade-offs and the capacity to predict environmental variation will likely influence the ultimate trajectory. Below we consider potential drivers and implications of this adaptive plasticity for eco-evolutionary dynamics as well as its applied relevance for the conservation and restoration of reefs.

### Adaptive phenotypic plasticity in coral

Phenotypic plasticity can facilitate acclimation over space and time by allowing organisms to match local phenotypic optima (6, 61). In reef-building corals, plastic responses in morphology (44, 62), bleaching (43), gene expression (52), and gene body methylation (63) alter colony phenotype in ways that are hypothesized to be beneficial, especially in the context of a stress response. In the absence of other performance data, however, it was unclear whether such individual plasticity would result in a net fitness benefit, particularly if significant costs were incurred by plastic individuals in non-stress contexts. Here, by explicitly linking morphological plasticity with growth and survival we show that an individual’s ability to be plastic yields a net positive fitness outcome in *A. cervicornis*. Specifically, increased plasticity was associated with both a higher average growth rate and an increased probability of survival (decreased mortality risk, Fig. 4A) strongly supporting an adaptive role.

Coral morphology is an important environmental interface as changes in size and shape can adjust the flow, temperature, and pH in and around a colony (64, 65) affecting both normal processes, such as nutrient uptake, as well as responses to thermal or acidification stress (66–68). In the Florida Keys, spatially or temporally variable conditions, such as temperature, select for local phenotypic optima (69, 70) that appear to promote the overall persistence of plastic *A. cervicornis* genotypes. Although sites did not appear to select for specific morphologies in terms of their clustering patterns, genets with the highest average survival occupied a broader morphospace (Fig. 4B), again supporting an adaptive role for morphological plasticity.

Coral meet many of the conditions predicted to favor the evolution of increased plasticity (14). Long generation times and high habitat heterogeneity across dispersal ranges expand the range of environments individuals experience within their lifetime (71, 72). Assuming a genetic basis to plasticity, positive selection in the form of increased survival and/or reproduction of more plastic individuals can enable genetic accommodation, making future generations more plastic (6). This evolution of plasticity may be modulated by strong selection events occurring in *A. cervicornis* and other reef building coral in the form of disease or bleaching events (40, 73), although the magnitude and direction of the effects will depend on the relationship between morphological plasticity and the response to stress. We did not detect any tradeoffs between plasticity, growth rate and survival in ambient conditions, but exploring additional costs or limits to morphological plasticity will be an important next step in understanding the future adaptive potential of plasticity, particularly in the face of new environmental extremes. Warming ocean temperatures are an ever-present threat to coral reefs and tradeoffs between plasticity and bleaching tolerance or recovery would severely limit the benefits of morphological plasticity as bleaching events become more frequent and severe (39, 40).

### Genotype-environment interactions limit survival and growth predictions

Survival and growth are two metrics commonly used to evaluate coral fitness because improved survival will positively impact population demographic rates (74–76) while faster growth can shorten the time to reproductive maturity (77). Although lifetime reproductive success is difficult to measure in annual broadcast spawning species such as coral, greater reproductive capacity has been observed in larger colonies (77). Significant variation in the Cox Proportional Hazard risk scores indicated that some genotypes (G41, G62, G13) were at greater risk of mortality than others. Similarly, reef sites also varied in their ability to support coral survival. These results align with previous findings of strong genotype and site effects on coral survival (32, 57) and suggest that certain genets (G36) and reef sites (E. Sambo) may be higher quality overall. When looking at the remaining genets and sites, however, genotype-by-site interactions for survival and lack of preservation in the rank order of genet survivorship between reef sites (Fig. 2C) indicate a limited ability to predict outcomes based on knowledge of genotypes or sites in isolation. This stands in contrast to a similar in situ transplant experiment by Drury et al. (33) that found site effects but no effect of genotype or the interaction on mortality. However, corals in this prior study endured a thermal bleaching event and variation in mortality was attributed to variation in bleaching among reef sites (33). While no bleaching was observed during the course of our experiment, our results indicate that under normal conditions, genet survivorship from a single site alone does not predict survival in other, even geographically neighboring, reefs.

Similarly, coral morphology appears to be influenced by a complex set of interacting factors that ultimately preclude identification of a globally top performing individual or ‘super coral’ (34). Genotype-by-site interactions were evident in absolute size which resulted in a different collection of genets representing the largest relative coral at each site after the 12 month outplant period (Fig. 3). This is the first evidence of GxE in *A. cervicornis* morphology, although this finding builds on earlier work in other branching coral species (53, 54, 78). It is important to note that statistical models included corals that experienced fragmentation, as this is an ecologically relevant phenomenon contributing to clonal reproduction (79). When we excluded fragmentation, significant genotype-by-site interactions disappeared. Similarly, (33) excluded negative growth and showed independent impacts of genotype and environment on TLE in *A. cervicornis*, but no GxE was detected. Significant fixed effects of genotype on both absolute size and growth rate for TLE, SA, V, and V_inter_ suggest that the intrinsic growth rate does vary among genets. However, this capacity may be limited by fragmentation, which also varies as a function of genotype and site. Ultimately however, population demographic and restoration success are based on size of coral colonies rather than their growth rate, indicating that GxE must be accounted for when developing conservation and restoration strategies.

Unlike earlier studies in both *A. cervicornis* (33) and its sister species, *A. palmata* (80), we find no effect of site on the majority of morphological and growth traits despite strong site effects on survival. Once thought of as noise, GxE allows for the presence of phenotypic variation among individuals in the absence of overall site effects and may provide a proximate explanation for their absence here. Alternatively, differences in experimental design may also play a role. Kuffer et al. (80) compared ramets outplanted over a large spatial scale (>300 km) compared to the ~60 km span covered in the present study. Similarly, Drury et al. (33) selected reef sites spanning inshore and offshore zones. Although reefs within the offshore zone vary in environmental conditions (56) larger differences are evident between inshore and offshore reefs in temperature, turbidity, and nutrients (56, 81) may have driven the pronounced site effects on morphology reported earlier (33). Finally, temporal variability in environmental stress can create periods of poor growth or high fragmentation followed by periods of recovery that may mask mean site effects and instead generate a site-by-time signal. We did observe significant site-by-time effects on the majority of morphological traits indicating that site effects are likely to be time dependent.

### The importance of fragmentation

Fragmentation is a vital part of the evolutionary ecology of *A. cervicornis* as it can significantly alter the demographic trajectory of populations through asexual propagation (79, 82), partial mortality, or death (83). In this experiment numerous instances of breakage had obvious negative impacts on colony size but were mostly nonfatal. This suggests that fragmentation can also impact growth beyond the immediate response to injury, such as increased productivity due to size-dependent growth rates that occur after size reduction (57). While sometimes considered random, fragmentation in this study occurred significantly more in some genets and sites. Differences in calcification rate among *A. cervicornis* genets (84, 85) mean that certain individuals can produce more dense skeletons faster, potentially making them less prone to breakage. Moreover, calcification is energetically expensive (86) and apparent trade-offs in skeletal density and colony size (84, 85) suggest different strategies for skeletal growth in this species that may lead to variation in the ability to withstand physical stress leading to breakage.

Spatial and temporal variation in hydrodynamic energy (87, 88), likely also imposes variable mechanical stress on coral colonies. Coral morphology may respond to these conditions (44, 89), but sudden or especially strong hydrodynamic forces are common sources of damage for branching corals in the Caribbean (90, 91). Human activity may also have contributed to fragmentation and anecdotally, higher tourist activity was observed at Looe Key and EDR. Although fragmentation was usually nonfatal in the focal ramet, we did not track the fate of newly generated fragments precluding determination of the ultimate effect on fitness. Regardless, the existence of non-random fragmentation reinforces the notion that accounting for fragmentation, rather than treating it as experimental error, will be important for accurately predicting changes in branching coral morphology and performance.

### Multivariate environmental conditions distinguish reefs

The offshore reef sites used here are in an area that has historically been treated as a single environmental unit (56). However, site specific variation in coral performance (Fig. 2B, Fig. 3) and in environmental condition (Fig. 1E) support the need for a more nuanced approach. Aggregate differences in abiotic conditions among sites with high average survival appear to be defined by high nitrogen concentrations, thermal predictability, light attenuation/turbidity, and low annual and average daily temperature ranges (Fig. S13-14). Similarly, the site with the lowest average survival, Bahia Honda, differentiated by historically high ranges of nitrite and total phosphorus concentrations, turbidity, and light attenuation (Fig. S14). This site also had the highest annual and daily temperature ranges, but lowest summer thermal predictability during the experimental period. Although no physical or chemical condition was individually correlated with mortality in the Bayesian models, fluctuating environmental conditions have been implicated in the conditioning of marine species to climate change (92, 93). While temperature variability can enhance coral tolerance (94, 95), the predictability of those fluctuations should also impact the ability to acclimate and adapt (96, 97). Thermal predictability was highest among the three sites with the highest survival (E. Sambo, Marker 32, and W. Sambo) and lowest at Bahia Honda yet no correlations were detected, suggesting the importance of a multivariate approach. The low temporal resolution of water quality metrics precluded obtaining similar measures of predictability in water chemistry during the experimental period. Future work quantifying environmental predictability in addition to fluctuations may yield additional insight into the conditions that support coral performance and plasticity.

Mean temperature was negatively associated with change in size of all morphological traits (Fig. 1F, S15-18), suggesting that cooler conditions promoted ramet growth. This is unsurprising considering the well documented negative impact of high temperatures on coral performance (98) and the fact that mean experimental temperatures were within or above the apparent optimum thermal range (ca 25-29°C) for *A. cervicornis* (99, 100). Interestingly, the number of days above 30.5°C seemed to encourage growth in V_inter_ but no other trait. Morphology has been shown to impact flow around a colony, altering heat flux at the coral surface (64, 68) with branching morphologies more capable of offloading heat compared to mounding coral (101). As *A. cervicornis* ramets become less compact by increasing the volume of their interstitial space (Fig. 1C), heat dissipation at the coral surface can also be expected to increase through a reduction in the thermal boundary layer (68, 101). While the offshore reef tract of the Florida Keys is typically considered environmentally contiguous, taken together these results suggest spatial variation in reef conditions independently and cumulatively shape coral performance.

### Conclusions

Current coral restoration strategies rely on transplanting clones across reefs varying in abiotic and biotic conditions (33, 37, 55) suggesting that plasticity will play a role in the success or failure of individual colonies. Adaptive morphological plasticity in *A. cervicornis* may enable genets to maintain fitness in response to changes in environmental conditions over time or space. Continued positive selection on intraspecific variation in plasticity, contingent on its freedom from tradeoffs, should promote the evolution of plasticity and therefore the acclimatory benefits associated with it. Environmental conditions can also promote or constrain the evolution of phenotypic plasticity (10, 96) and while there appears to be sufficient variation within the *A. cervicornis* habitat range to induce plasticity at present, its relative benefits will likely also be dependent on the ability to predict environmental fluctuations, which may prove challenging in the face of continued climate change.

## Materials and Methods

### Experimental design

Ten coral genets maintained long-term (5+ years) at Mote Marine Laboratory’s in situ coral nursery (Table S10) were outplanted in a multi-site transplant study under FKMNS permits 2015-163-A1 and 2018-035. In April 2018, 270 coral (mean TLE of 8.4 cm) ramets representing 10 genets (27 ramets per genet) affixed to concrete pucks were photographed following Million et al. (102) and manually measured for TLE immediately before transplantation to nine active restoration sites (Table S1, Fig. 1). Three ramets per genet (n=270 fragments) were randomly outplanted at each site with one ramet allocated to each of three ten-coral arrays (Fig. 1A). Coral pucks were attached to bare reef substrate using marine epoxy over 4 days, from April 21 to April 25.

Outplant sites were resurveyed in July 2018, October 2018, January 2019, and April 2019. Ramets were individually re-photographed and measured by-hand at the first four time-points for TLE. Survivorship was recorded during site surveys and later confirmed with photographs. Breakage was recorded via the photographic time series and through negative growth measures in the resulting trait dataset (Supplemental Methods). Ramets where both the coral tissue and ceramic puck were missing indicated technical failure of the marine epoxy rather than a true biological loss and were excluded from subsequent analyses.

### Phenotyping

Photographs taken in situ were used to generate individual 3D models of each coral ramet in Metashape 1.5.4 (Agisoft LLC, St. Petersburg, Russia) using a high-throughput pipeline (102). Specifications for model building and all scripts can be found at https://github.com/wyattmillion/Coral3DPhotogram. 3D models were imported into Meshlab v2020.6 (103) to measure four growth-related traits following protocols described in Million et al. (102) and detailed in the Supplementary Methods: TLE, SA, V, and V_inter_. We assessed the final shape of colonies that survived to T12 with no detectable fragmentation by calculating SA-to-V and TLE-to-V ratios, in addition to packing, convexity, and sphericity (59). Among this subset, genets were more equally represented (n=5-12 ramets per genets) than sites (n=2-18 ramets per site). These traits were used in a principal components analysis to determine how ramets clustered in morphospace as a function of genet and site.

### Environmental data

All reef sites are located along the offshore reef tract of the Lower Florida Keys at a depth of 5.6m to 9.1m. HOBO Pendant Temperature loggers (Onset Computer Corp.; Bourne, MA), set to record hourly, were attached to the reef substrate directly adjacent to the outplanted corals, and exchanged with new loggers on subsequent site visits. Hourly temperature records were used to calculate annual mean, annual range, average daily range, maximum monthly mean, days and hours above 30.5°C or 32°C, and summer thermal predictability. Thermal predictability was quantified for July through September only as highly variable temperatures at or above the bleaching threshold of 30.5°C are expected during this window (104). Predictability was calculated as the sum of autocorrelation over a series of lags until autocorrelation reached zero, i. e. the point at which current temperatures are no longer informative of future conditions (105, 106). Quarterly concentrations of benthic nitrite, nitrate, ammonia, dissolved organic and inorganic nitrogen, soluble reactive phosphorus, total phosphorus, total nitrogen, N:P ratio, silicate, dissolved oxygen, total organic carbon, as well as turbidity and light attenuation for each outplant site were obtained from the Southeast Environmental Research Center (SERC) water quality dataset (Florida International University) associated with each site (Fig. 1A, serc.fiu.edu/wqmnetwork/FKNMS-CD/DataDL.htm).

### Statistical analysis

All statistical analyses were performed in R version 3.6.3 (107) and scripts can be found at github.com/wyattmillion/Acer_Morphological_Plasticity. Cox Proportional Hazard models were fitted to outplant survival data using the coxme (108) and survival (109) packages. Consistency in genet rank order across sites was quantified with Pearson’s correlations. Cumulative Linked Mixed Models assessing the ordinal response of fragmentation were implemented with the package ordinal (110) to test for effects of genotype and outplant site on cumulative breakage events summed within a ramet over time. Fisher Exact Tests were used to determine enrichment of fragmentation among *A. cervicornis* size classes.

Effects of genotype, site, time, all associated interactions, and initial size on colony morphology (~size) and growth rate in TLE, SA, V, and V_inter_ were tested with linear mixed effects models implemented with the package lmer (111). Fragmentation and outplant array nested within-site were included as random effects. When calculating growth rate, ramets experiencing fragmentation (evidenced by a negative growth rate for a ramet over a 3-month interval) were removed from the dataset and replaced with NA for the time-point in which fragmentation occurred.

The extent of plasticity was quantified across each time interval using the joint regression framework (58, 112, 113) where genet mean sizes at each site were regressed against the site-wide mean. Regression coefficients representing plasticity in TLE, SA, V, and V_inter_ were correlated with Cox mortality risk scores and average genet growth rates using Pearson Correlations.

SERC water quality parameters and thermal characteristics were used to describe environmental variation among reef sites. We calculated both the overall mean and the range of benthic nutrient concentrations, turbidity, and light attenuation over the entire length of the SERC dataset (Spring 1995 to Spring 2019) and over the experimental period (April 2018 to 2019). The historical and contemporary SERC data were used to identify differences between reef sites along with high resolution temperature data collected during the experimental period. An analysis of variance was used to identify significant differences among sites for independent parameters. A principal components analysis was used to explain variation among sites using all parameters simultaneously.

Bayesian negative binomial generalized linear models implemented in R2jags (114) were used to test for the impact of environmental parameters on the growth and survival of ramets at reef sites. Model power was improved by using size, risk score, and environmental data over each of the four time intervals to increase sample size and by removing highly correlated environmental variables.

## Acknowledgements

We are grateful to Yingqi Zhang, Hunter Ramo, Cory Walter, and Joseph Kuehl for their help photographing coral in situ. Eric Million helped design and build equipment for in situ 3D photogrammetry. Phoebe Chang contributed to image preprocessing and Alexandra Stella and Aryana Volk helped with model validation. Elaina Graham generously facilitated access to the Heidelberg Lab high performance computer. Training in Bayesian analyses from Dr. Robert van Woesik was facilitated by the NSF Coral Bleaching RCN Early Career Training Program. Figure 1 was created with BioRender.com. All fieldwork was conducted under permits FKNMS-2015-163-A1 and FKNMS-2018-035.

## Author Contributions

CDK and CJK designed the experiment and obtained funding; WCM, MR, EB, CJK and CDK conducted the experiment; All authors collected data; WCM and CDK performed data analysis; WCM and CDK wrote the initial manuscript while all authors revised and contributed to the final version.

## Conflict of Interest

The authors declare no competing interest.

## Funding Information

This research was supported by National Oceanic and Atmospheric Administration Coral Reef Conservation Program grant NA17NOS4820084 and private funding from the Alfred P. Sloan and Rose Hills Foundations.

## Data Accessibility

The datasets and scripts used in this study can be found at https://github.com/wyattmillion/Acer_Morphological_Plasticity.

## Notes

https://github.com/wyattmillion/Acer_Morphological_Plasticity

### Competing Interest Statement

The authors have declared no competing interest.

