## SupplementaryMaterial for "Evidence for adaptive morphological plasticity in the Caribbean coral, *Acropora cervicornis*"

**Supplemental Methods**

Ten coral genets maintained long-term (5+ years) at the Mote Marine Lab Summerland Key *in situ* coral nursery (Table S1 for coordinates) were selected for use in a multi-site transplant study in partnership with Mote Marine Lab’s Staghorn Restoration Program under FKMNS permits 2015-163-A1 and 2018-035. On March 19, 2018, 300 coral (mean TLE of 8.4 cm) ramets representing 10 genets (27 ramets per genet) were transferred from hanging trees, fixed to ceramic pucks with marine epoxy and attached to benthic structures in preparation for outplanting. In April 2018, each ramet was photographed for 3D photogrammetry following the protocol described in Million et al. 2021 and manually measured for total linear extension (TLE) immediately before transplantation to nine established restoration sites (Table S1 for coordinates) in the Lower Florida Keys. Sites were chosen based on their overlap with actively permitted *A. cervicornis* restoration activities and proximity to permanent Southeast Environmental Research Center (SERC) Water Quality Monitoring stations (SERC, Florida International University). Three ramets per genet (n=270 fragments) were randomly outplanted at each site with one ramet allocated to each of three ten-coral arrays. Arrays were spaced approximately 1-m apart and the subsite locations were selected based on substrate type, prioritizing hard bottom. Coral pucks were attached to bare reef substrate using marine epoxy leaving approximately 10 cm between each puck. Outplantating occurred over 4 days due to weather and time constraints, from April 21 to April 25.

Outplant sites were resurveyed every three months for a total of four time-points throughout a one-year period, in July 2018, October 2018, January 2019, and April 2019. Ramets were individually photographed for 3D photogrammetry-based reconstruction of growth metrics and total linear extension was measured by-hand for the first four time-points. Survivorship was recorded during site surveys and confirmed with photographs post-hoc. Ramets were recorded as alive if any amount of living tissue was visible. Breakage was recorded both via the photographic time series and through negative growth measures in the resulting trait dataset. Ramets where both the coral tissue and ceramic puck were missing were discarded from the analyses as we attribute these losses to technical failure of the marine epoxy upon outplanting rather than true biological mortality.

*Phenotyping*

Photographs taken *in situ* were used to generate individual 3D models of each coral ramet in Metashape 1.5.4 (Agisoft LLC, St. Petersburg, Russia) using the high-throughput pipeline described in Million et al. (1). Models were built on Dell PowerEdge R910 with an Intel® Xeon® Processor E7-4850 with Metashape manually limited to 20-40 CPUs and 250 GB of RAM. Specifications for model building and all scripts can be found at <https://github.com/wyattmillion/Coral3DPhotogram>. 3D models in Wavefront format (.obj) from Metashape were imported into Meshlab v2020.6 (2) to measure four growth-related traits following protocols described in Million et al. (1): total linear extension (TLE, Johnson et al. 2011), surface area (SA), volume (V), and volume of the convex hull. The volume of the convex hull represents the smallest sized, convex mesh that encompasses all points of an object and is used to calculate the volume of interstitial space (V_inter_) for each coral colony by subtracting the volume of the colony from the volume of the convex hull.

*Environmental data*

HOBO Pendant Temperature loggers (Onset Computer Corp., Bourne, MA) were affixed to the reef substrate with plastic zip ties and set to record hourly for six month periods and were redeployed on subsequent site visits. SERC has monitored water quality quarterly at a network of 270 sites throughout south Florida and the Florida Keys beginning in 1995 (3). Concentrations of benthic nitrite, nitrate, ammonia, dissolved organic and inorganic nitrogen, soluble reactive phosphorus, total phosphorus, total nitrogen, N:P ratio, silicate, dissolved oxygen, total organic carbon, turbidity, and light attenuation for each monitoring station associated with an outplant site were obtained from the publicly available SERC database (SERC, Florida International University). Depth of reef sites range from 5.6m to 9.1m.

*Statistical analysis*

All statistical analyses were performed in R version 3.6.3 (4). Raw data and scripts can be found at github.com/wyattmillion/Acer_Morphological_Plasticity. Cox Proportional Hazard models were fitted to outplant survival data using *coxme* *(5)*and *survival* *(6)* packages in order to test for effects of genotype, environment, and genotype-by-environment interactions. Effects of fragmentation were included in mixed effects Cox models by accounting for the cumulative number of breakage events experienced by each ramet. When models including the interaction of genotype and outplant site did not converge, an additive model was used to reliably estimate coefficients for fixed effects. Consistency in genet rank, ordered 1 to 10 by increasing mortality risk, across sites was quantified with Pearson’s correlations. Cumulative Linked Mixed Models were applied with the package *ordinal* *(7)* to test for effects of genotype and outplant site on the ordinal response variable of cumulative breakage events summed within a ramet across time. Effects of size preceding the presence/absence of fragmentation was tested for using a binomial logistic regression.

Effects of genotype, environment, and genotype-by-environment interactions on colony morphology (~size) and growth rate were tested with linear mixed effects models in the package *lmer* *(8)**.* Morphological change was assessed in two ways for each trait. Both the absolute size and the monthly growth rate (determined for each 3-month time interval) were calculated for TLE, SA, V, and V_inter_. Trait values were either square-root or log(x+n) transformed to meet assumptions of normality and account for zero-values. Models for absolute size included genotype, outplant site, time point, all two and three-way interactions and initial size as fixed effects. Fragmentation and outplant array nested within site were included as random effects. To accurately quantify growth rates, ramets experiencing fragmentation (evidenced by a negative growth rate for a ramet over a 3-month interval) were removed from the dataset and replaced with NA for only the time-point in which fragmentation occurred. With this reduced data set, we assessed the fixed effects of genotype, site, time point, all associated interactions, and ramet size at the start of each 3-month interval on growth rate in each trait, including a random effect of array nested within site.

Correlations between the extent of plasticity and hazard ratio (mortality risk) for each genet were used to address the adaptive role of phenotypic plasticity. The extent of plasticity was quantified using the joint regression framework(9, 10) in which genet mean trait values in each site are regressed against the environmental mean across all genets at each site. The resulting coefficient gives genet plasticity relative to the population; a coefficient >1 indicates enhanced plastic response in a genet relative to the population while a coefficient <1 suggests less responsiveness. Plasticity measured through joint regression, as compared to traditional two-environment approaches (11), incorporate data over multiple environments and generate a unitless metric comparable across traits. Coefficients representing plasticity in TLE, surface area, volume, and volume of interstitial space in addition to trait means were used as covariates in a linear model to explain variation in genet hazard ratios estimated from Cox models. This process is commonly used to estimate selection on plasticity in plant systems (9, 12, 13) as it controls for covariance between independent variables.

SERC water quality parameters and thermal characteristics were used to describe environmental variation among reef sites. We calculated both the mean and the average annual range of benthic μM concentrations of nitrite, nitrate, ammonia, total nitrogen, total phosphorus, total organic nitrogen, total organic carbon, silicate, dissolved oxygen, as well as turbidity and light attenuation over the entire extent of the SERC dataset (Spring 1995 to Spring 2019) and over the experimental period (April 2018 to 2019). The historical and contemporary SERC data were used to identify differences between reef sites along with high resolution temperature data collected during the experimental period. Annual mean, annual range, average daily range, maximum monthly mean, days and hours above 30.5°C or 32°C, and thermal predictability for each site was generated from the hourly temperature data. Thermal predictability was quantified for only the Summer months (June through September) because this period represents highly variable temperatures at or above the bleaching threshold of 30.5°C (14). Autocorrelation was calculated with the *acf* function of the *stats* package over a series of lags until autocorrelation reached zero. The sum of autocorrelation coefficients in that series of lags was then used to summarize the predictability of each site. Sites with larger sums are expected to be more predictable as current temperatures are more informative of temperatures longer into the future. An analysis of variance was used to identify significant differences among sites for independent parameters. A principal components analysis was used to explain variation among sites using all parameters simultaneously.

Bayesian negative binomial generalized linear models implemented in *R2jags* (15) were used to simultaneously test for the impact of multiple environmental parameters on the growth and survival of coral colonies at outplant sites. To improve model fit, change in size and risk score were calculated for each time interval and paired with environmental parameters over the same interval. For SERC data, a single data point represented each interval while thermal conditions were calculated within each interval. Highly correlated environmental parameters, identified with Pearson correlations, were removed from the analysis leaving 11 metrics to be used in the Bayesian model: total phosphorus, ammonia, total organic carbon, dissolved oxygen, as well as light attenuation, turbidity, days above 30.5°C, thermal predictability, average daily range, seasonal range, mean temperature.

**Supplemental Figures and Tables**

Supplementary Table 1: Outplant sites and SERC water quality monitoring station coordinates

| Outplant sites | | | SERC water quality monitoring stations | | |
| --- | --- | --- | --- | --- | --- |
| Site | Lon | Lat | Station | Lon | Lat |
| Bahia Honda | -81.24203 | 24.58908 | 256 (Bahia Honda) | -81.2341 | 24.585 |
| Big Pine Shoals | -81.32655 | 24.56867 | 259 (Big Pine) | -81.3217 | 24.5704 |
| Looe Key | -81.40214 | 24.54667 | 263 (Looe Key) | -81.3974 | 24.5484 |
| Dave's Ledge | -81.48734 | 24.53051 | 267 (Dave’s Ledge) | -81.4873 | 24.5301 |
| Maryland Shoals | -81.56991 | 24.51036 | 270 (Maryland Shoals) | -81.5643 | 24.5216 |
| E. Sambo | -81.65961 | 24.49305 | 273 (E. Sambo) | -81.6571 | 24.4929 |
| W. Sambo | -81.70334 | 24.48268 | 403 (W. Sambo) | -81.7 | 24.483 |
| Marker 32 | -81.7426 | 24.47408 | 276 (Marker 32) | -81.7374 | 24.4754 |
| Eastern Dry Rocks | -81.84412 | 24.45948 | 280 (EDR) | -81.8437 | 24.4536 |

Supplementary Table 2: Results of Cox proportional hazard ratio models for genets

|  | Coef | exp(coef) | se(coef) | z | p |
| --- | --- | --- | --- | --- | --- |
| Genotype1 | 0.9524501 | 2.592053 | 1.228389 | 0.78 | 0.44 |
| Genotype50 | 1.2405797 | 3.457617 | 1.155948 | 1.07 | 0.28 |
| Genotype3 | 1.1978105 | 3.312855 | 1.158254 | 1.03 | 0.3 |
| Genotype44 | 1.0984207 | 2.999425 | 1.155609 | 0.95 | 0.34 |
| Genotype7 | 1.5670542 | 4.792509 | 1.120372 | 1.4 | 0.16 |
| Genotype31 | 1.6818513 | 5.375498 | 1.099799 | 1.53 | 0.13 |
| Genotype13 | 2.3255781 | 10.232594 | 1.063958 | 2.19 | 0.029 |
| Genotype62 | 2.5519734 | 12.832402 | 1.059458 | 2.41 | 0.016 |
| Genotype41 | 2.6081417 | 13.573804 | 1.052236 | 2.48 | 0.01 |

Supplementary Table 3: Results of Cox proportional hazard ratio models for sites

| Site | Coef | exp(coef | se(coef) | z | p |
| --- | --- | --- | --- | --- | --- |
| Marker 32 | 1.0708734 | 2.917927 | 1.232165 | 0.87 | 0.38 |
| W. Sambo | 1.3784374 | 3.968695 | 1.118639 | 1.23 | 0.22 |
| Big Pine | 1.3089321 | 3.702218 | 1.119653 | 1.17 | 0.24 |
| Dave's Ledge | 1.4134808 | 4.110237 | 1.120982 | 1.26 | 0.21 |
| Looe Key | 2.1242445 | 8.366574 | 1.083228 | 1.96 | 0.05 |
| EDR | 2.0674828 | 7.904900 | 1.082746 | 1.91 | 0.056 |
| Maryland Shoals | 2.1523159 | 8.604763 | 1.056446 | 2.04 | 0.042 |
| Bahia Honda | 2.5649031 | 12.999399 | 1.045958 | 2.45 | 0.014 |

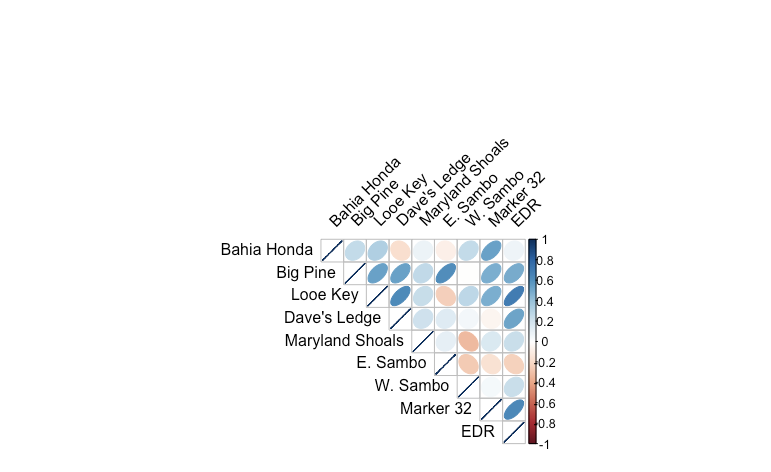

Supplementary Figure 1: Pairwise genet rank correlations across outplant sites ordered by geographic location (longitudinally). Shape and color of ellipses are proportional to the strength and direction of correlation.

Supplementary Table 4: Results of cumulative linked model testing the effects of genet and site on ramet fragmentation

| Sites | | | | | Genotypes | | | | |
| --- | --- | --- | --- | --- | --- | --- | --- | --- | --- |
|  | Estimate | Std. Error | z value | Pr(>\|z\|) |  | Estimate | Std. Error | z value | Pr(>\|z\|) |
| E. Sambo | 0.5077 | 0.3317 | 1.531 | 0.12584 | 36 | 0.4645 | 0.3358 | 1.383 | 0.16652 |
| W. Sambo | 0.7426 | 0.3273 | 2.268 | 0.02330 | 50 | 0.2953 | 0.3379 | 0.874 | 0.38227 |
| Big Pine | 0.8121 | 0.3339 | 2.432 | 0.01502 | 3 | 0.6699 | 0.3364 | 1.991 | 0.04644 |
| Dave's Ledge | 0.9349 | 0.3287 | 2.844 | 0.00445 | 44 | 0.6699 | 0.3383 | 1.980 | 0.04766 |
| Looe Key | 0.8999 | 0.3257 | 2.763 | 0.00573 | 7 | 0.5157 | 0.3436 | 1.501 | 0.13344 |
| EDR | 0.6931 | 0.3358 | 2.064 | 0.03902 | 31 | 0.6021 | 0.3347 | 1.799 | 0.07206 |
| Maryland Shoals | 1.3802 | 0.3270 | 4.221 | 2.44e-05 | 13 | 0.7520 | 0.3365 | 2.235 | 0.02544 |
| Bahia Honda | 1.5764 | 0.3291 | 4.79 | 1.67e-06 | 62 | 0.2679 | 0.3400 | 0.788 | 0.43075 |
|  |  |  |  |  | 41 | 0.5835 | 0.3369 | 1.732 | 0.08329 |

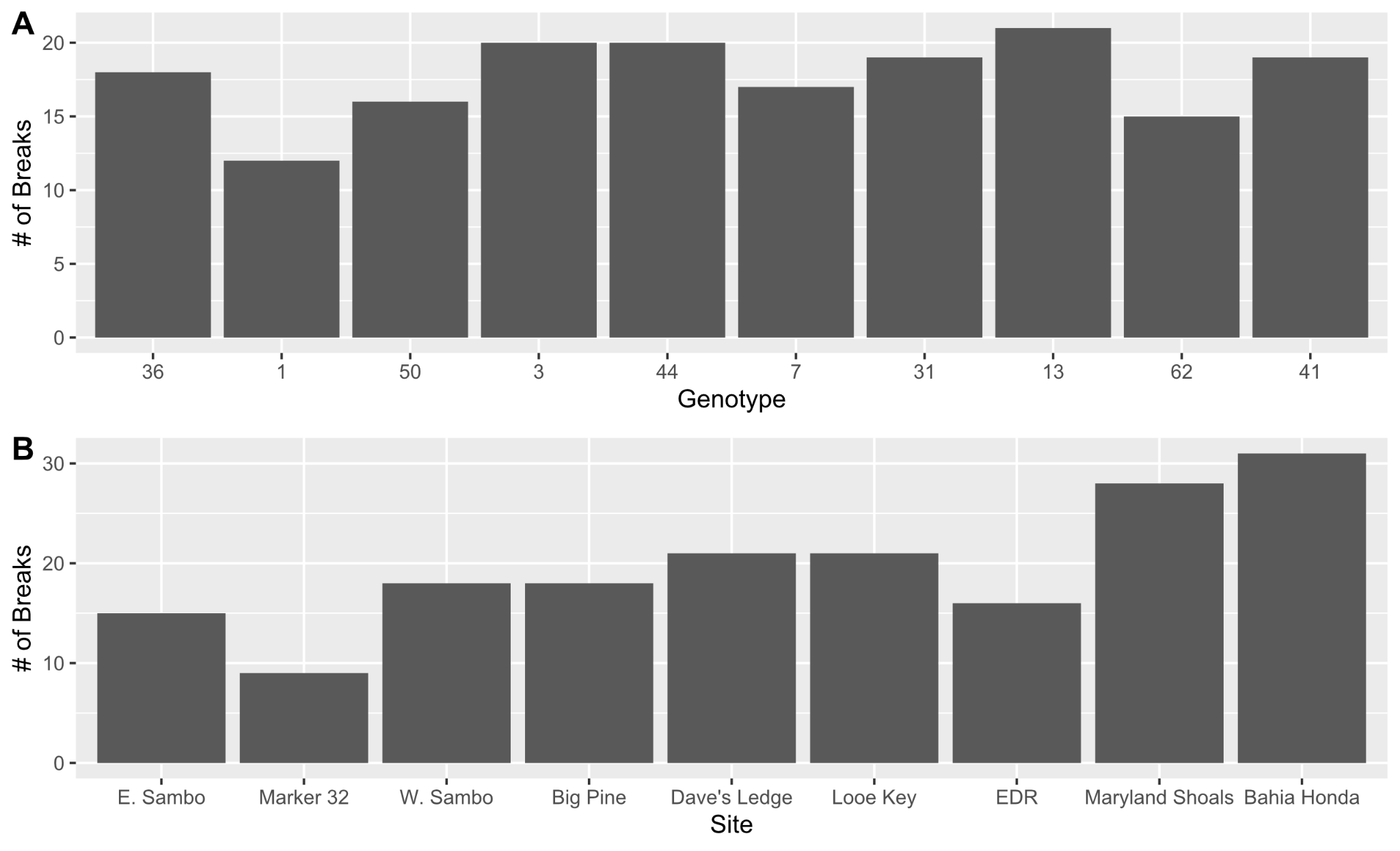

Supplementary Figure 2: Total number of observed fragmentation events for genets (A) and sites (B) across the one-year experimental period.

Supplementary Table 5: Summary of results for Fisher Exact test of the prevalence of detectable fragmentation in *A. cervicornis* size classes.

| Size Class | n | p | p.adj p. |
| --- | --- | --- | --- |
| >5 cm | 897 | 1.56E-6 | 9.36E-6 |
| 5-10 cm | 897 | 5.52E-5 | 2.76E-4 |
| 10-15 cm | 897 | 1 | 1 |
| 15-20 cm | 897 | 0.485 | 1 |
| 20-30 cm | 897 | 1 | 1 |
| 30+ cm | 897 | 0.358 | 1 |

Supplementary Table 6: Summary of fixed effects for absolute size in TLE, SA, V and V_inter_

| Trait | Factor | Sum Sq | Mean Sq | Num DF | DenDF | F value | Pr(>F) |
| --- | --- | --- | --- | --- | --- | --- | --- |
| SA | **Genotype** | 228.76 | 25.42 | 9 | 482.92 | 5.2064 | 9.06E-07 |
| TLE | **Genotype** | 30.545 | 3.3939 | 9 | 496.4 | 6.4518 | 1.07E-08 |
| V | **Genotype** | 32.874 | 3.6526 | 9 | 481.46 | 9.5383 | 1.98E-13 |
| Vinter | **Genotype** | 68.554 | 7.617 | 9 | 467.44 | 6.6733 | 5.23E-09 |
| SA | **Genotype:Site** | 839.91 | 11.83 | 71 | 479.29 | 2.4231 | 1.82E-08 |
| TLE | **Genotype:Site** | 81.196 | 1.1436 | 71 | 492.65 | 2.174 | 8.25E-07 |
| V | **Genotype:Site** | 70.096 | 0.9873 | 71 | 478.13 | 2.5781 | 1.53E-09 |
| Vinter | **Genotype:Site** | 202.854 | 2.857 | 71 | 463.97 | 2.5031 | 5.74E-09 |
| SA | Genotype:Site:time | 594.05 | 2.87 | 207 | 475.75 | 0.5879 | 0.999992 |
| TLE | Genotype:Site:time | 64.679 | 0.311 | 208 | 488.71 | 0.5911 | 1 |
| V | Genotype:Site:time | 47.526 | 0.2296 | 207 | 474.72 | 0.5996 | 0.999985 |
| Vinter | Genotype:Site:time | 116.492 | 0.571 | 204 | 460.7 | 0.5003 | 1 |
| SA | Genotype:time | 164.55 | 6.09 | 27 | 476.13 | 1.2484 | 0.183772 |
| TLE | Genotype:time | 9.829 | 0.364 | 27 | 488.91 | 0.692 | 0.8771 |
| V | Genotype:time | 10.524 | 0.3898 | 27 | 475.08 | 1.0179 | 0.441626 |
| Vinter | Genotype:time | 13.982 | 0.518 | 27 | 461.02 | 0.4537 | 0.9925 |
| SA | Site | 74.99 | 9.37 | 8 | 18.05 | 1.9198 | 1.19E-01 |
| TLE | Site | 3.813 | 0.4766 | 8 | 17.99 | 0.9059 | 5.33E-01 |
| V | Site | 7.544 | 0.943 | 8 | 18.09 | 2.4624 | 5.33E-02 |
| Vinter | Site | 12.019 | 1.502 | 8 | 18.24 | 1.316 | 2.96E-01 |
| SA | **Site:time** | 219.69 | 9.15 | 24 | 477.38 | 1.875 | 0.007664 |
| TLE | Site:time | 13.699 | 0.5708 | 24 | 490.21 | 1.0851 | 3.56E-01 |
| V | **Site:time** | 18.814 | 0.7839 | 24 | 476.4 | 2.0471 | 2.65E-03 |
| Vinter | Site:time | 37.087 | 1.545 | 24 | 463.34 | 1.3538 | 1.24E-01 |
| SA | **T0_SA** | 436.82 | 436.82 | 1 | 483.12 | 89.4773 | < 2.2e-16 |
| TLE | **T0_TLE** | 17.872 | 17.872 | 1 | 475.02 | 33.9751 | 1.03E-08 |
| V | **T0_V** | 17.317 | 17.3167 | 1 | 491.95 | 45.2208 | 4.90E-11 |
| Vinter | **T0_Vinter** | 55.842 | 55.842 | 1 | 468.85 | 48.9231 | 9.23E-12 |
| SA | **time** | 1236.44 | 412.15 | 3 | 477.54 | 84.4239 | < 2.2e-16 |
| TLE | **time** | 83.662 | 27.8874 | 3 | 490.2 | 53.0146 | < 2.2e-16 |
| V | **time** | 64.458 | 21.4858 | 3 | 476.57 | 56.1081 | < 2.2e-16 |
| Vinter | **time** | 161.364 | 53.788 | 3 | 464.72 | 47.1238 | < 2.2e-16 |

Supplementary Table 7: Summary of random effects on absolute size in TLE, SA, V, and V_inter_

| trait |  | npar | logLik | AIC | LRT | Df | Pr(>Chisq) |
| --- | --- | --- | --- | --- | --- | --- | --- |
| **TLE** | **(1 \| Site:Array)** | 354 | -799.21 | 2306.4 | 26.338 | 1 | 2.87E-07 |
| **TLE** | **(1 \| CulumativeBreaks)** | 354 | -853.03 | 2414.1 | 133.988 | 1 | < 2.2e-16 |
| **SA** | **(1 \| Site:Array)** | 353 | -1338.9 | 3383.9 | 33.072 | 1 | 8.88E-09 |
| **SA** | **(1 \| CulumativeBreaks)** | 353 | -1383.7 | 3473.3 | 122.483 | 1 | < 2.2e-16 |
| **V** | **(1 \| Site:Array)** | 353 | -705.6 | 2117.2 | 29.383 | 1 | 5.94E-08 |
| **V** | **(1 \| CulumativeBreaks)** | 353 | -760.71 | 2227.4 | 139.605 | 1 | < 2.2e-16 |
| **Vinter** | **(1 \| Site:Array)** | 350 | -951.64 | 2603.3 | 39.209 | 1 | 3.81E-10 |
| **Vinter** | **(1 \| CulumativeBreaks)** | 350 | -983.4 | 2666.8 | 102.73 | 1 | < 2.2e-16 |

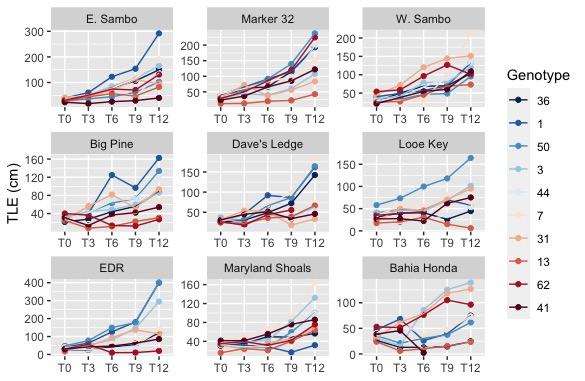

Supplementary Figure 3: Growth curves in SA. Genet averages at each site are colored in decreasing overall survival from blue to red.

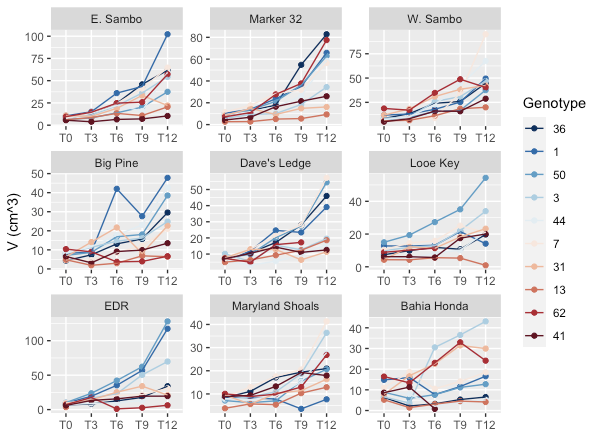

Supplementary Figure 4: Growth curves in V. Genet averages at each site are colored in decreasing overall survival from blue to red.

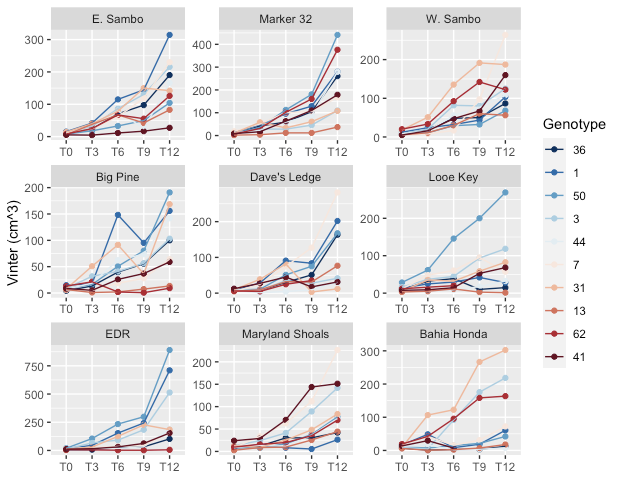

Supplementary Figure 5: Growth curves in V_inter_. Genet averages at each site are colored in decreasing overall survival from blue to red.

Supplementary Table 8: Summary of linear fixed effects for growth rate in TLE, SA, V and V_inter_

| Trait | Factor | Sum Sq | Mean Sq | Num DF | DenDF | F value | Pr(>F) |
| --- | --- | --- | --- | --- | --- | --- | --- |
| SA | **Genotype** | 14.294 | 1.588 | 9 | 331.93 | 1.6839 | 0.091632 |
| TLE | **Genotype** | 6.502 | 0.7224 | 9 | 341.01 | 4.0994 | 5.09E-05 |
| V | **Genotype** | 6.618 | 0.7354 | 9 | 319.51 | 2.8875 | 0.002701 |
| Vinter | **Genotype** | 4.655 | 0.5173 | 9 | 314.95 | 2.0695 | 0.031896 |
| SA | Genotype:Site | 70.673 | 1.024 | 69 | 291.44 | 1.0813 | 0.324806 |
| TLE | Genotype:Site | 15.82 | 0.2228 | 71 | 305.87 | 1.2597 | 0.0958539 |
| V | Genotype:Site | 19.679 | 0.2852 | 69 | 307.21 | 1.1196 | 0.259268 |
| Vinter | Genotype:Site | 23.119 | 0.3303 | 70 | 276.28 | 1.3167 | 0.063247 |
| SA | Genotype:Site:time | 144.302 | 0.756 | 191 | 310.69 | 0.8015 | 9.53E-01 |
| TLE | Genotype:Site:time | 34.243 | 0.1793 | 191 | 328.49 | 1.0182 | 4.40E-01 |
| V | Genotype:Site:time | 35.751 | 0.1902 | 188 | 301.91 | 0.7468 | 9.85E-01 |
| Vinter | Genotype:Site:time | 44.74 | 0.2472 | 181 | 297.03 | 0.9894 | 5.28E-01 |
| SA | **Genotype:time** | 55.574 | 2.058 | 27 | 332.49 | 2.1831 | 0.000797 |
| TLE | **Genotype:time** | 8.858 | 0.3281 | 27 | 348.14 | 1.8629 | 0.0065125 |
| V | Genotype:time | 9.541 | 0.3534 | 27 | 313.22 | 1.3875 | 0.099349 |
| Vinter | **Genotype:time** | 13.152 | 0.4871 | 27 | 314.72 | 1.9495 | 0.003913 |
| SA | Site | 13.433 | 1.679 | 8 | 15.21 | 1.7799 | 0.158868 |
| TLE | Site | 2.102 | 0.2627 | 8 | 16.01 | 1.4912 | 0.2358322 |
| V | **Site** | 6.258 | 0.7823 | 8 | 15.46 | 3.0699 | 0.028047 |
| Vinter | Site | 2.162 | 0.2702 | 8 | 16.81 | 1.0809 | 0.421249 |
| SA | Site:time | 32.699 | 1.362 | 24 | 333.21 | 1.445 | 8.35E-02 |
| TLE | **Site:time** | 10.609 | 0.442 | 24 | 349.66 | 2.5098 | 1.53E-04 |
| V | **Site:time** | 10.827 | 0.4511 | 24 | 314.67 | 1.7714 | 1.56E-02 |
| Vinter | **Site:time** | 11.752 | 0.4897 | 24 | 316.85 | 1.9594 | 0.005308 |
| SA | **size** | 91.326 | 91.326 | 1 | 26.58 | 96.8685 | 2.35E-10 |
| TLE | **size** | 15.476 | 15.4763 | 1 | 20.61 | 87.8828 | 7.03E-09 |
| V | **size** | 11.356 | 11.356 | 1 | 302.72 | 44.592 | 1.16E-10 |
| Vinter | **size** | 26.01 | 26.0104 | 1 | 72.21 | 104.099 | 1.21E-15 |
| SA | **time** | 44.401 | 14.8 | 3 | 240.44 | 15.6703 | 2.43E-09 |
| TLE | **time** | 6.556 | 2.1855 | 3 | 228.84 | 12.3819 | 1.56E-07 |
| V | **time** | 9.445 | 3.1485 | 3 | 318.51 | 12.3629 | 1.14E-07 |
| Vinter | **time** | 10.58 | 3.5268 | 3 | 257.45 | 14.0975 | 1.55E-08 |

Supplementary Table 9: Summary of random effects on growth rate in TLE, SA, V, and V_inte_

| Trait | Factor | npar | logLik | AIC | LRT | Df | Pr(>Chisq) |
| --- | --- | --- | --- | --- | --- | --- | --- |
| TLE | (1 \| CulumativeBreaks) | 337 | -361.08 | 1396.2 | 0 | 1 | 1 |
| TLE | (1 \| Site:Array) | 337 | -363.41 | 1400.8 | 4.6555 | 1 | 0.03095 |
| SA | (1 \| CulumativeBreaks) | 335 | -641.38 | 1952.8 | 0 | 1 | 1 |
| SA | (1 \| Site:Array) | 335 | -643 | 1956 | 3.2458 | 1 | 0.07161 |
| V | (1 \| CulumativeBreaks) | 332 | -395.19 | 1454.4 | 2.5096 | 1 | 0.1132 |
| V | (1 \| Site:Array) | 332 | -394.92 | 1453.8 | 1.9622 | 1 | 0.1613 |
| Vinter | (1 \| CulumativeBreaks) | 326 | -390.51 | 1433 | 0.0041 | 1 | 0.9487 |
| Vinter | (1 \| Site:Array) | 326 | -398.15 | 1448.3 | 15.2826 | 1 | 9.26E-05 |

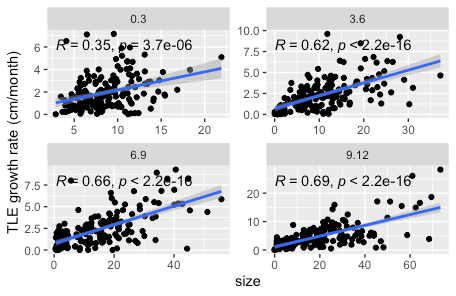

Supplementary Figure 6: Relationship of growth rate in total linear extension (cm/month) and ramet size at each time interval: Apr -Jul 2018 (0.3), Jul -Oct (3.6), Oct 2018-Jan 2019 (6.9), Jan-Apr 2019 (9.12)

Supplementary Figure 7: Relationship between growth rate in total linear extension (cm/month) standardized to existing ramet biomass and ramet size at each time interval: Apr-Jul 2018 (0.3), Jul-Oct (3.6), Oct 2018-Jan 2019 (6.9), Jan-Apr 2019 (9.12)
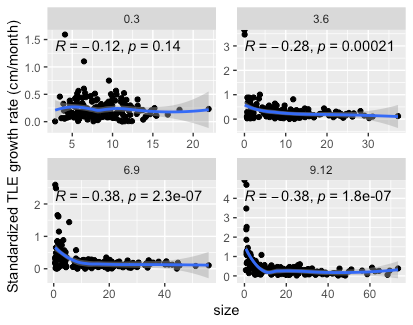

Supplementary Figure 8: Pearson correlations between genet plasticity calculated with a joint regression at a given time point and genet risk score calculated across the entire experimental period. Rows of plots show plasticity-risk score relationships in a trait at the four time intervals while columns. Black line and shaded region show line of best fit and 95% confidence interval, respectively.
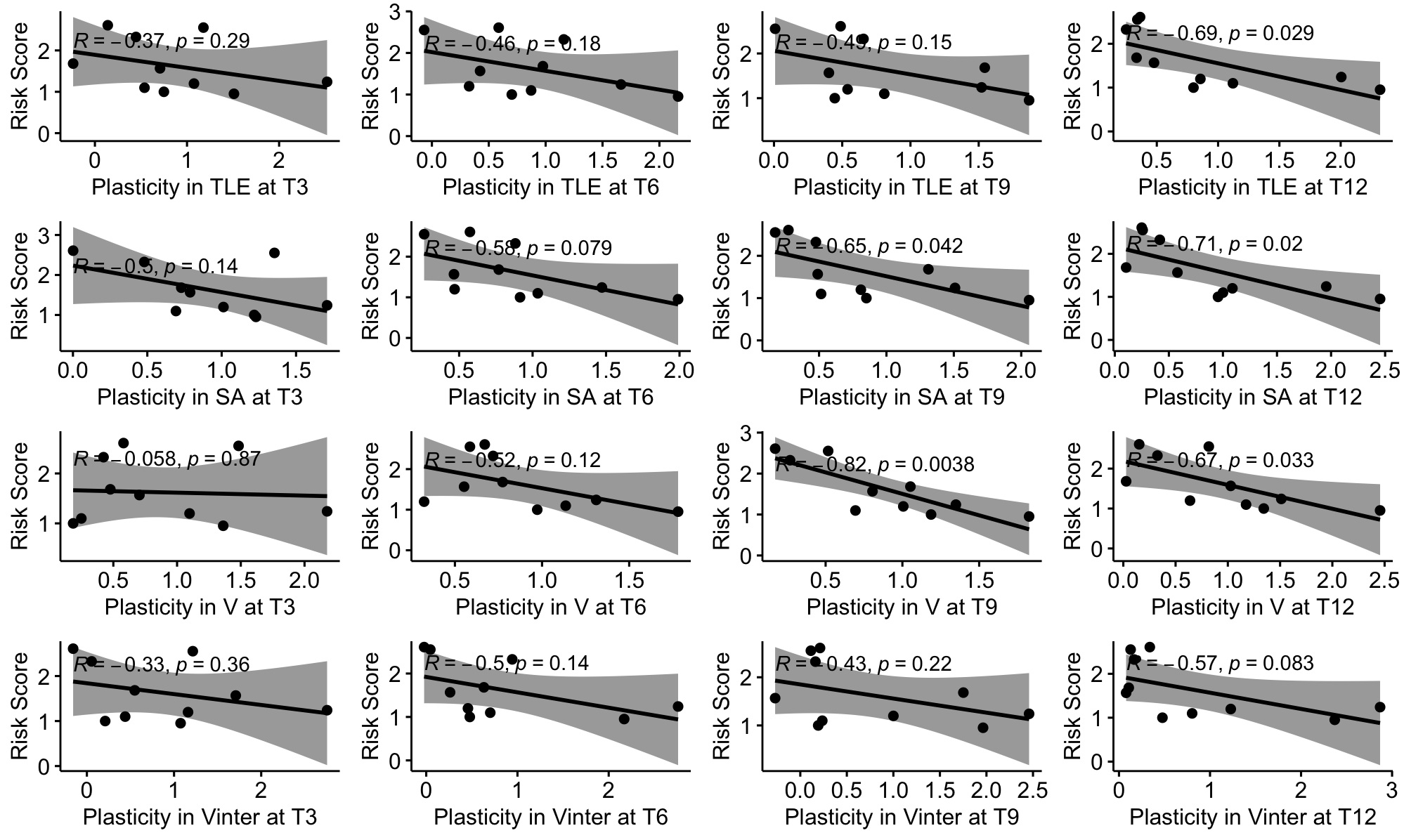

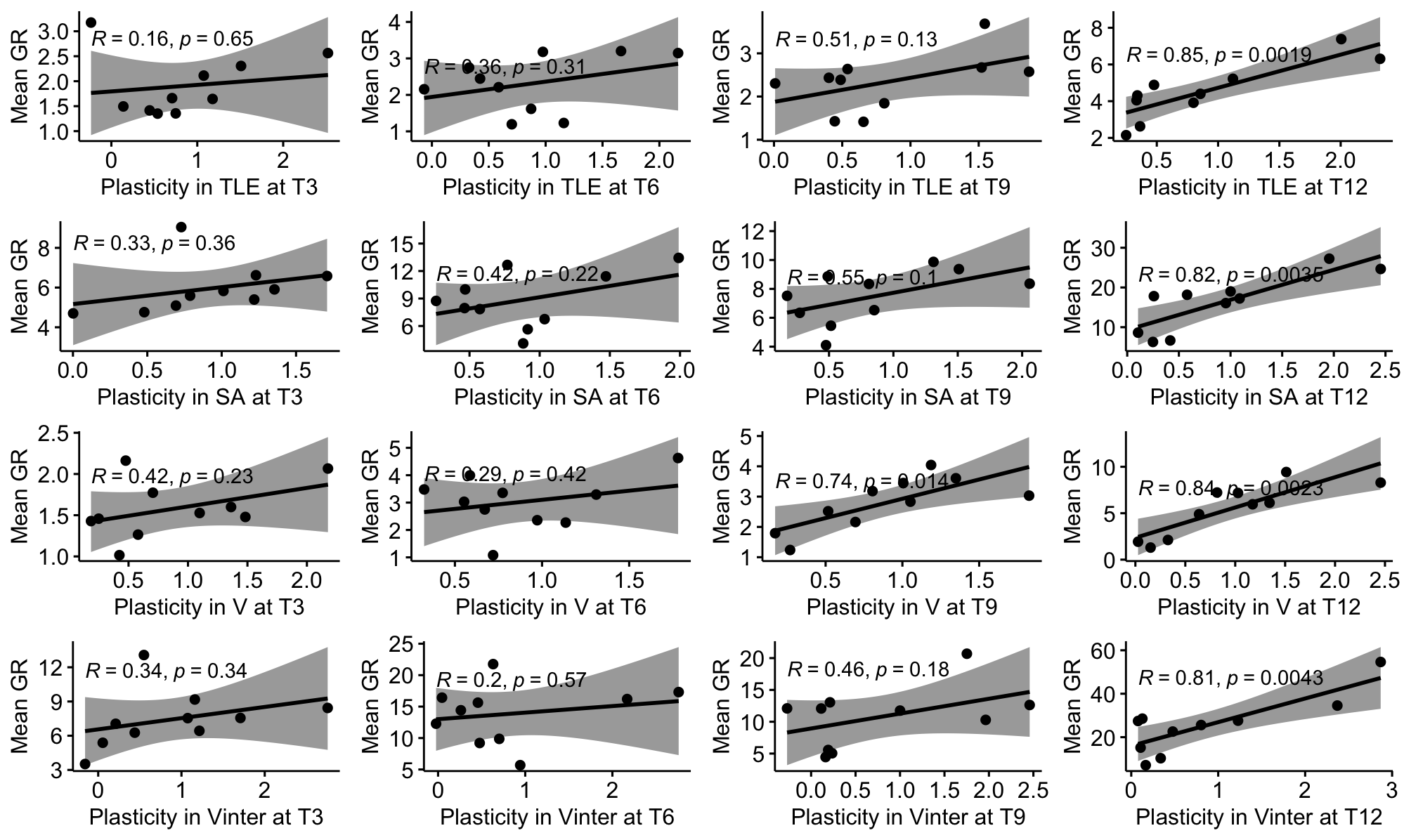

Supplementary Figure 9: Pearson correlations between genet plasticity calculated with a joint regression at a given time point and genet average growth rate (unit/month) at a given time point. Rows of plots show plasticity-growth rate relationships in a trait at the four time intervals while columns. Black line and shaded region show line of best fit and 95% confidence interval, respectively.

Supplementary Figure 10: Pearson correlations between genet average growth rate (unit/month) at a given time point and genet risk score calculated across the entire experimental period. Rows of plots show growth rate-risk score relationships in a trait at the four time intervals while columns. Black line and shaded region show line of best fit and 95% confidence interval, respectively.
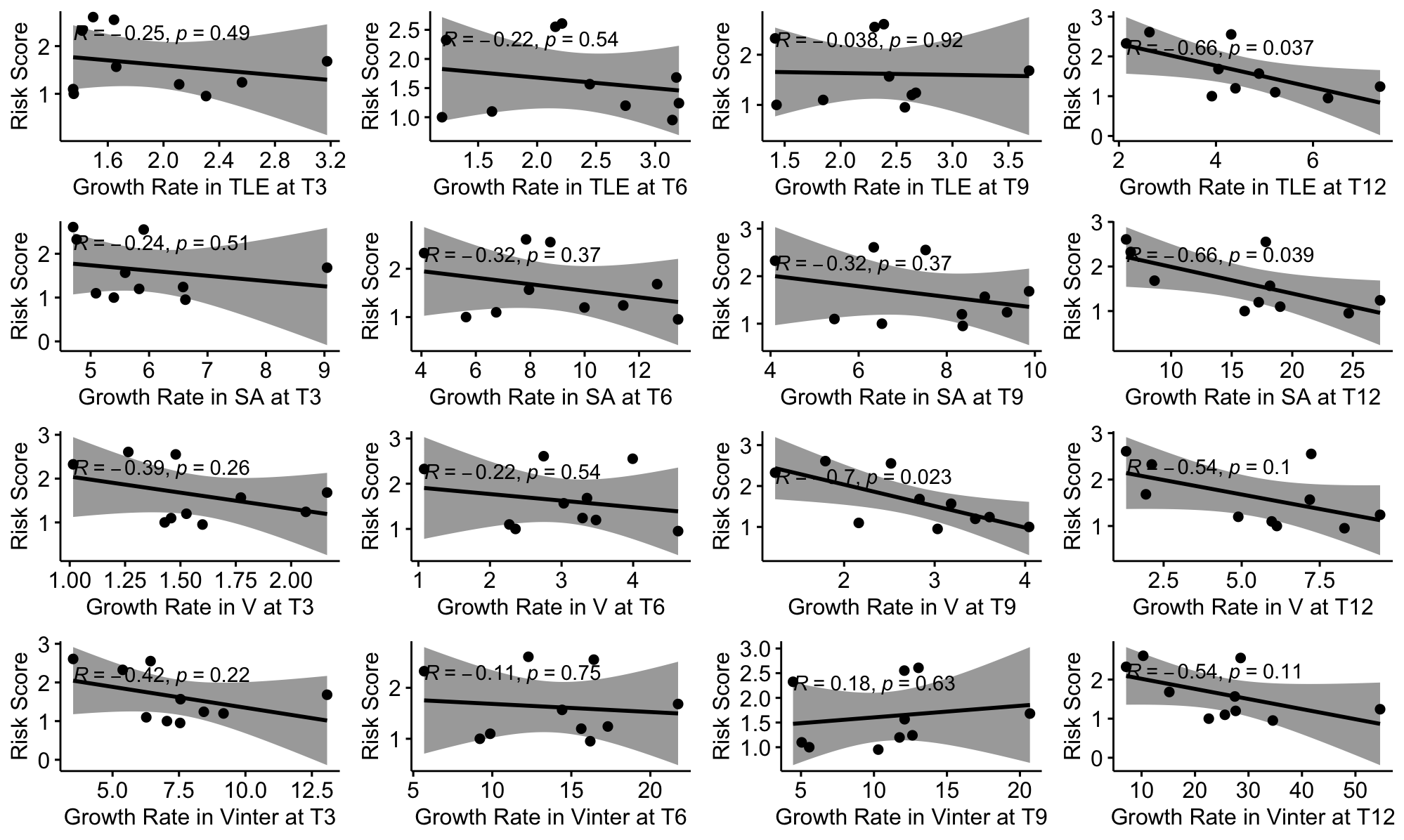

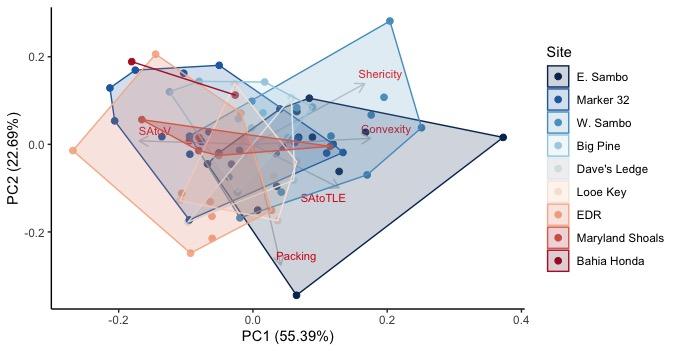

Supplemental Figure 11: Principal components analysis of ramet invariant morphology measured with SA-to-V ratio, TLE-to-V ratio, packing, convexity, and sphericity. Points represent ramets surviving to T12 without experiencing fragmentation (n=81) color by sites decreasing in survival from blue to red. Polygons define the PCA space occupied by ramets of each site.

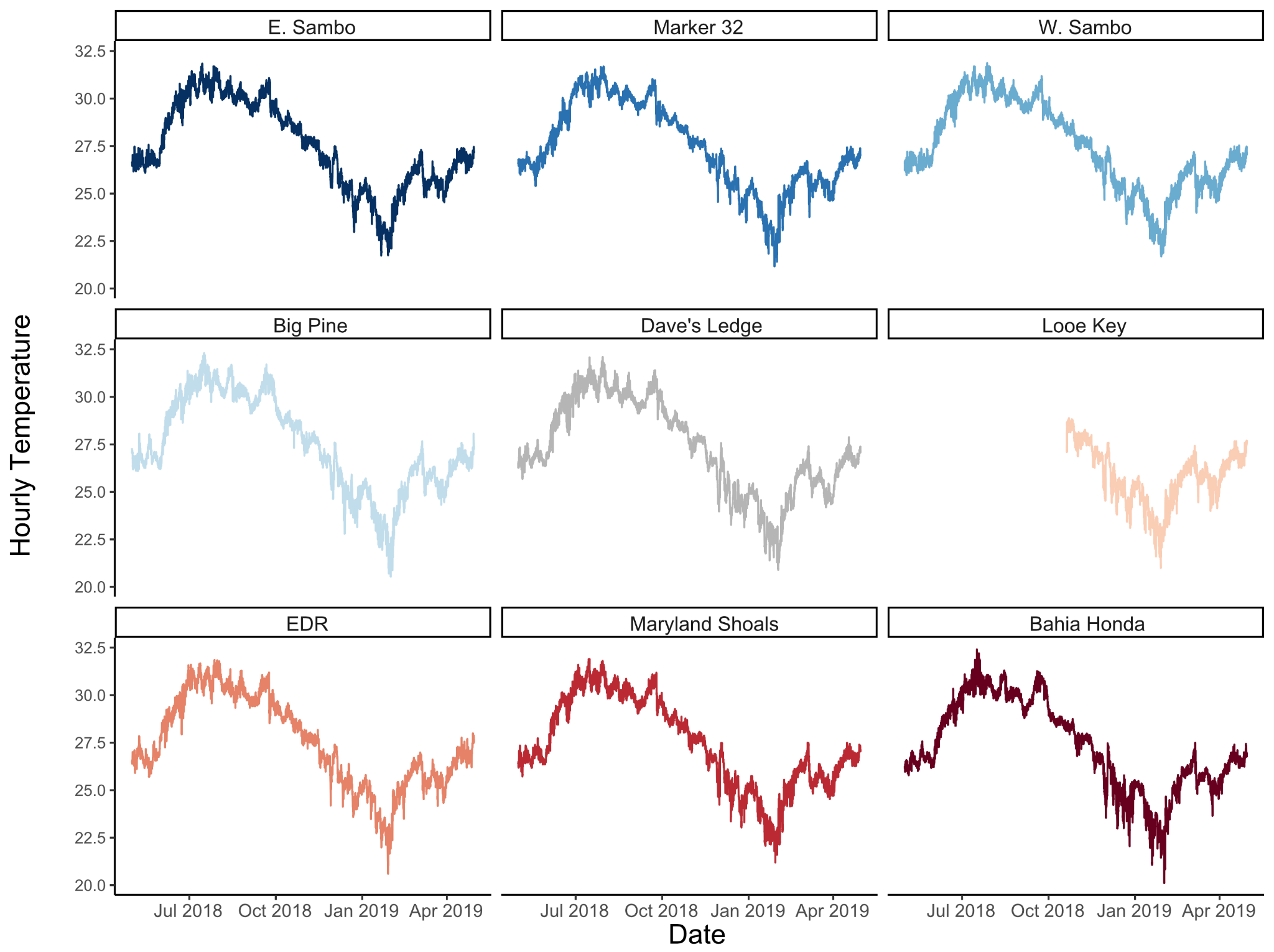

Supplementary Figure 12: Hourly temperature data from April 2018 to April 2019 for each outplant reef site.

Supplementary Table 10: Maximum monthly mean, annual mean, annual range, average daily range, number of days where temperatures reached at least 30.5C (TDD30.5) or 32C (TDD32), number of hours where temperatures were at or above 30.5C (TDH30.5) or 32C (TDH32), and Summer thermal predictability for outplant reef sites calculated over the one-year experimental period.

| Site | Max Monthly Mean | Annual Mean | Annual Range | Avg Daily Range | TDD30.5 | TDD32 | TDH  30.5 | TDH32 | Predictability |
| --- | --- | --- | --- | --- | --- | --- | --- | --- | --- |
| E. Sambo | 30.833 | 27.474 | 10.12 | 0.628 | 57 | 0 | 824 | 0 | 209.141 |
| Marker 32 | 30.774 | 27.386 | 10.489 | 0.587 | 54 | 0 | 824 | 0 | 195.195 |
| W. Sambo | 30.810 | 27.448 | 10.216 | 0.658 | 54 | 0 | 806 | 0 | 214.138 |
| Big Pine | 30.963 | 27.450 | 11.772 | 0.694 | 69 | 3 | 1154 | 27 | 134.525 |
| Dave's Ledge | 30.803 | 27.305 | 11.185 | 0.636 | 57 | 2 | 870 | 4 | 138.193 |
| Looe Key* | 28.215 | 25.809 | 7.857 | 0.790 | 0 | 0 | 0 | 0 | NA |
| EDR | 30.882 | 27.489 | 11.265 | 0.650 | 67 | 0 | 1060 | 0 | 186.321 |
| Maryland Shoals | 30.855 | 27.408 | 10.693 | 0.700 | 63 | 0 | 863 | 0 | 182.886 |
| Bahia Honda | 30.653 | 27.276 | 12.256 | 0.706 | 49 | 3 | 781 | 10 | 105.005 |

Supplementary Table 11: Summary of results for analysis of variance for SERC water quality parameters using the full historical dataset (Spring 1995 -Spring 2019).

| Parameter | DF | Sum Sq | Mean Sq | F value | Pr (>F) |
| --- | --- | --- | --- | --- | --- |
| Nitrite | 8 | 0.0117 | .00147 | 1.517 | 0.147 |
| Nitrate | 8 | 1.12 | 0.141 | 2.405 | 0.0145 |
| Ammonia | 8 | 1.23 | 0.1543 | 1.027 | 0.414 |
| Total Nitrogen | 8 | 450 | 56.28 | 0.813 | 0.591 |
| Total Organic Nitrogen | 8 | 437 | 54.65 | 0.77 | 0.63 |
| Total Phosphorus | 8 | 0.017 | 0.00215 | 0.203 | 0.99 |
| Soluble Reactive Phosphorus | 8 | 0.0051 | 0.00064 | 0.576 | 0.798 |
| Turbidity | 8 | 21.7 | 2.718 | 0.694 | 0.697 |
| Total Organic Carbon | 8 | 54788 | 6849 | 0.934 | 0.488 |
| Silicon Dioxide | 8 | 16.9 | 2.112 | 2.03 | 0.0405 |
| Salinity | 8 | 0.4 | 0.04944 | 0.329 | 0.955 |
| Dissolved Oxygen | 8 | 0.7 | 0.0819 | 0.138 | 0.997 |
| Light Attenuation | 8 | 0.27 | 0.0342 | 0.734 | 0.661 |
| Oxygen Saturation | 8 | 127 | 15.88 | 0.139 | 0.997 |

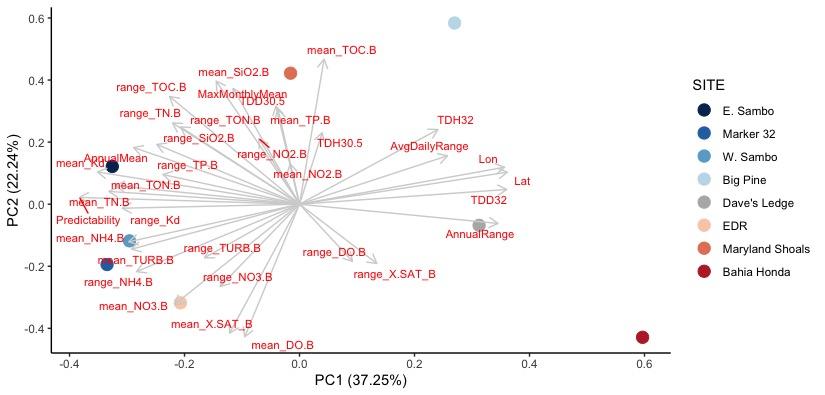

Supplementary Figure 13: Principal components analysis of contemporary (April 2018-April 2019) SERC water quality data and thermal characteristics for outplant sites used. Grey arrows show vector loadings with labels at each of the tips in red. Sites are colored by decreasing survival from blue to red. Looe Key is removed due to a lack of temperature data from April to October 2018.

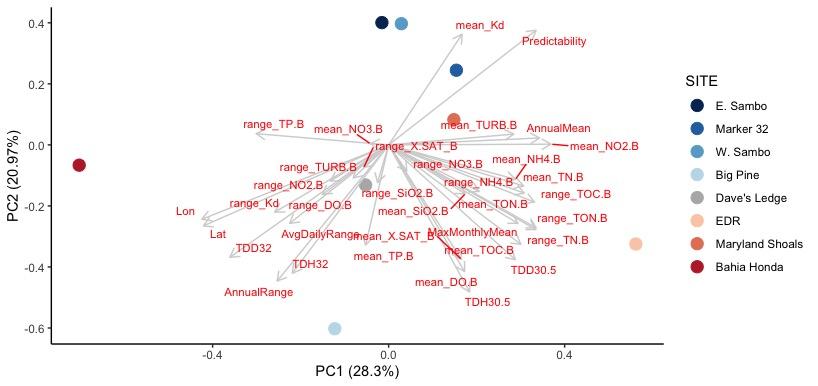

Supplementary Figure 14: Principal components analysis of historical (April 2018-April 2019) SERC water quality data and thermal characteristics for outplant sites used. Grey arrows show vector loadings with labels at each of the tips in red. Sites are colored by decreasing survival from blue to red. Looe Key is removed due to a lack of temperature data from April to October 2018.

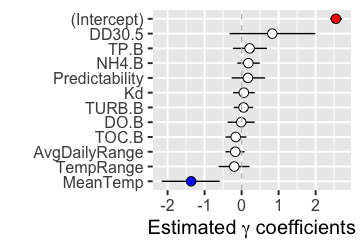

Supplementary Figure 15: Output from the Bayesian negative binomial generalized linear model for total linear extension. Change in average TLE over a given time interval was the response variable for environmental predictors measured over the respective time interval. Points represent mean effect while error bars represent 95% credible intervals. Points are colored blue (negative effect on response variable), red (positive effect) if the credible interval does not overlap zero, and white otherwise.

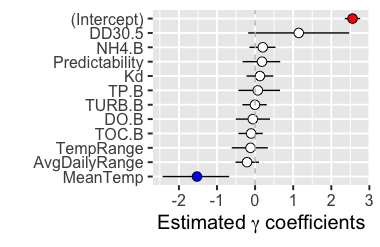

Supplementary Figure 16: Output from the Bayesian negative binomial generalized linear model for surface area. Change in average SA over a given time interval was the response variable for environmental predictors measured over the respective time interval. Points represent mean effect while error bars represent 95% credible intervals. Points are colored blue (negative effect on response variable), red (positive effect) if the credible interval does not overlap zero, and white otherwise.

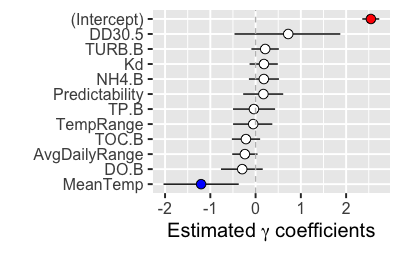

Supplementary Figure 17: Output from the Bayesian negative binomial generalized linear model for volume. Change in average V over a given time interval was the response variable for environmental predictors measured over the respective time interval. Points represent mean effect while error bars represent 95% credible intervals. Points are colored blue (negative effect on response variable), red (positive effect) if the credible interval does not overlap zero, and white otherwise.

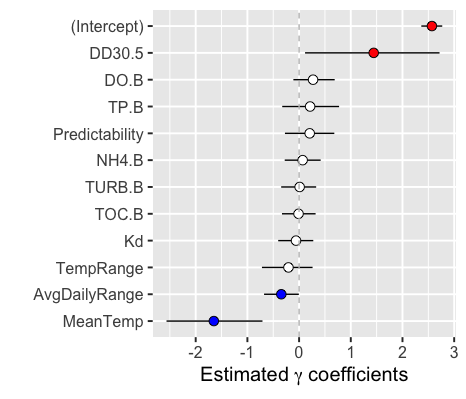

Supplementary Figure 18: Output from the Bayesian negative binomial generalized linear model for volume of interstitial space. Change in average V_inter_ over a given time interval was the response variable for environmental predictors measured over the respective time interval. Points represent mean effect while error bars represent 95% credible intervals. Points are colored blue (negative effect on response variable), red (positive effect) if the credible interval does not overlap zero, and white otherwise.

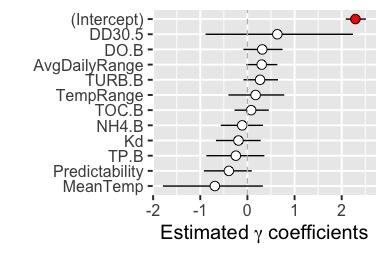

Supplementary Figure 19: Output from the Bayesian negative binomial generalized linear model for risk score. Site risk score over a given time interval was the response variable for environmental predictors measured over the respective time interval. Points represent mean effect while error bars show 95% credible intervals. Points are colored blue (negative effect on response variable), red (positive effect) if the credible interval does not overlap zero, and white otherwise.

1.	W. C. Million, S. O’Donnell, E. Bartels, C. D. Kenkel, Colony-Level 3D Photogrammetry Reveals That Total Linear Extension and Initial Growth Do Not Scale With Complex Morphological Growth in the Branching Coral, Acropora cervicornis. *Frontiers in Marine Science* **8**, 384 (2021).

2.	P. Cignoni, *et al.*, *MeshLab: an Open-Source Mesh Processing Tool* (2008).

3.	H. O. Briceño, J. N. Boyer, 2018 Annual Report of the Water Quality Monitoring Project for the Water Quality Protection Program of the Florida Keys National Marine Sanctuary (2019) (January 26, 2022).

4.	R Core Team, *R: A language and environment for statistical computing* (2020).

5.	T. M. Therneau, *coxme: Mixed Effects Cox Models* (Comprehensive R Archive Network (CRAN), 2020).

6.	T. M. Therneau, T. Lumley, E. Atkinso, C. Crowson, *survival: Survival Analysis* (Comprehensive R Archive Network (CRAN), 2022) (March 3, 2022).

7.	R. H. B. Christensen, *Regression Models for Ordinal Data* (2019).

15.	Y.-S. Su, M. Yajima, R2jags: Using R to Run “JAGS.” *Comprehensive R Archive Network (CRAN)* (2021) (March 2, 2022).
